# Trafficking of Cholesterol from Lipid Droplets to Mitochondria in Bovine Luteal Cells: Acute Control of Progesterone Synthesis^1^

**DOI:** 10.1101/409599

**Authors:** Michele R. Plewes, Crystal Cordes, Emilia Przgrodzka, Heather Talbott, Jennifer Wood, Andrea Cupp, John S. Davis

**Author notes:** **Correspondence:** Olson Center for Women’s Health, University of Nebraska Medical Center, 983255 Nebraska Medical Center, Omaha, NE 68198-3255, USA.

## Abstract

The corpus luteum (CL) is a transient endocrine gland that synthesizes and secretes the steroid hormone, progesterone. Progesterone biosynthesis is a complex process, converting cholesterol via a series of enzymatic reactions, into progesterone. Lipid droplets in luteal cells store cholesterol in the form of cholesterol esters, which can be utilized for steroidogenesis. In small luteal cells, luteinizing hormone (LH) increases intracellular cAMP concentrations leading to activation of protein kinase A (PKA), which phosphorylates downstream proteins, such as hormone sensitive lipase (HSL). Phosphorylation of HSL at Ser563 leads to increased HSL activation and association with lipid droplets, events which theoretically release cholesterol, which can be used for progesterone synthesis. Bovine CL were obtained from a local abattoir, dispersed, and luteal cells were enriched for SLC via centrifugal elutriation. Our results reveal that LH, forskolin, and cAMP induce HSL phosphorylation at Ser563and Ser660. Moreover, inhibiting HSL activity attenuates LH-induced P4 synthesis. Confocal analysis revealed that LH stimulates translocation of HSL to lipid droplets and mitochondria. Furthermore, LH increased trafficking of cholesterol from the lipid droplets to the mitochondria which was dependent on both PKA and HSL activation. These results demonstrate cholesterol stored in lipid droplets are utilized for LH-induced progesterone biosynthesis. Likewise, PKA-induced activation of HSL is required for release and trafficking of cholesterol from the lipid droplets to the mitochondria. Taken together, these findings support a role for a PKA/HSL signaling pathway in response to LH and demonstrate the dynamic relationship between PKA, HSL, and the lipid droplets in the synthesis of progesterone.

**Highlights:**

- LH and PKA induce HSL phosphorylation at Ser563and Ser660
- HSL is required for optimal LH-induced P4 synthesis
- LH stimulates translocation of HSL to lipid droplets and mitochondria
- LH stimulated trafficking of cholesterol from lipid droplets to mitochondria

## Introduction

The corpus luteum (CL) is an ovarian endocrine gland that synthesizes and secretes the steroid hormone, progesterone, which is essential for the establishment and maintenance of pregnancy. Late in the follicular phase, the anterior pituitary gland synthesizes and secretes the gonadotropin, luteinizing hormone (LH), which is responsible for the follicular rupture and release of the ovum [1-3], referred to as ovulation. In addition to triggering ovulation, LH plays a critical role in the development of the CL and stimulation of progesterone production. Following ovulation, LH causes the theca and granulosa cells of the ovulated follicle to differentiate into small and large steroidogenic luteal cells, respectively [4-6]. Luteinizing hormone also promotes progesterone biosynthesis in luteal cells. Luteinizing hormone stimulates adenylyl cyclase leading to increases in intracellular cAMP [7], activation of PKA, and ultimately stimulation of steroidogenesis [8, 9]. Both the small and large luteal cells respond to LH stimulation, however, the small luteal cells are many times more responsive to LH and activators of the cAMP/protein kinase A (PKA) signaling pathway [10, 11] when compared to large luteal cells.

Progesterone biosynthesis is a complex process, converting cholesterol via a series of enzymatic reactions, into progesterone. Intracellular cholesterol is transported across the outer mitochondria membrane to the inner membrane using the steroidogenic acute regulatory protein (STAR) [12]. Cholesterol side-chain cleavage enzyme (CYP11A1), located within the inner membrane of the mitochondria, mediates the initial enzymatic reaction, converting cholesterol to pregnenolone [13], which exits the mitochondria and is converted to progesterone by the enzyme 3β-hydroxysteroid dehydrogenase (3BHSD) located in the endoplasmic reticulum [14]. Luteal cells are unique from other steroidogenic tissues in that they contains copious amounts of the steroidogenic enzyme required for optimal progesterone biosynthesis without changes in gene expression. Studies have demonstratred the conversion of pregnenolone to progesterone does not appear to limit progesterone biosythesis [15, 16]. Therefore, a key step in progesterone biosynthesis is cholesterol avalibility. Although steroidogenic cells can synthesize cholesterol de novo, majority of the cholesterol in luteal cells comes from the blood in the form of lipoproteins (LDL and/or HDL). Lipoproteins are internalized either through receptor-mediated endocytic or selective cellular uptake [17], where cholesterol is sorted from lipoproteins within endosomes. Endosomal cholesterol is then trafficked to the mitochondrion for progesterone biosynthesis or stored as cholesterol esters in lipid droplets [14, 18]. In small luteal cells, the acute actions of LH on steroidogenesis require maximum cholesterol availability for optimal progesterone production.

Lipid droplets are unique organelles within the cell that serve as lipid reservoirs and are abundant in both small and large luteal cells. Unlike adipose tissue and other ovarian cells (theca and granulosa cells), the lipid droplets present in luteal cells are enriched with cholesterol esters, which may be utilized for steroidogenesis. However, cholesterol esters stored in lipid droplets must first be hydrolyzed at the 3^rd^ carbon of the cholesterol ring, to be utilized for progesterone biosynthesis. Hormone-sensitive lipase (HSL) is an intracellular neutral lipase that is expressed in wide variety of tissues, including adipocytes [19, 20], macrophages [21, 22], adrenal glands [23, 24], liver [25], testis [26, 27], and the ovaries [28], and displays a broad substrate specificity, hydrolyzing triacylglycerol, diacylglycerol, monoacylglycerol, retinyl esters, and cholesterol esters [29].

Different from other lipases, HSL activity is highly regulated by phosphorylation/dephosphorylation. Phosphorylation of HSL occurs at multiple sites, including Ser660 and Ser563, which are believed to stimulate catalytic activity specific for triacylglycerol and cholesteryl esters [30-32]. In steroidogenic tissues, hormonal activation of PKA, stimulates the phosphorylation of HSL at Ser563 [33-35], which may be responsible for the neutral cholesteryl ester hydrolase activity that is observed in these tissues. Direct evidence has demonstrated HSL knockout mice, lack neutral cholesteryl ester hydrolase activity in both the adrenal or testis for those animals lacking HSL [26]. Another study observed, HSL knockout mice having decreased steroid hormone production leading to infertility. This advocates the possibility that HSL may provide the unesterified, free-cholesterol available for rapid progesterone production in luteal cells.

It is well documented that HSL hydrolyzes cholesterol esters in steroidogenic cells and is critically involved in the intracellular processing and availability of cholesterol. Furthermore, luteal cells are abundant in lipid droplets enriched with cholesterol esters. It is believed that through its action as a neutral cholesteryl ester hydrolase, HSL is involved in regulating intracellular cholesterol metabolism and, thus, contributing to a variety of pathways in which cells utilize cholesterol. We hypothesize that LH-induced cAMP/PKA signaling pathways regulate HSL activity, hydrolyzing cholesterol esters stored in lipid droplets for steroidogenesis in luteal cells. The objectives of the current study were to determine 1) the effect of LH and PKA signaling on phosphoylation of HSL, 2) the influence of HSL on LH-induced P4 production, 3)the effects of LH on colocalization of HSL with lipid droplets and mitochondria, 4) whether cholesterol esters stored in luteal lipid droplets are ultized for LH-induced progestrone production, and 5) the influence of PKA and HSL on trafficking of cholesterol from lipid droplet-derived cholesterol esters to the mitochondria.

## Methods and Materials

### Reagents

Penicillin G-sodium, streptomycin sulfate, HEPES, bovine serum albumin (BSA), Deoxyribonuclease l, fetal bovine serum (FBS), Tris-HCl, sodium chloride, ethylenediaminetetraacetic acid (*EDTA),* ethylene *glycol*-bis(β-aminoethyl ether)-N,N,N’,N’-*tetraacetic* acid (EGTA), sodium fluoride, Na4O2O7, Na3VO4, Triton X-100, Glycerol, dodecyl sodium sulfate, β-mercaptoethanol, bromophenol blue, Tween-20, paraformaldehyde, and acetone were all purchased from Sigma-Aldrich (St. Louis, MO, USA). The phosphate buffer solution, DMEM (Calcium-free, 4.0 g/L glucose), Penicillin Streptomycin Solution, trypan blue, Mitochondrial extraction kit, and Lipofectamine RNAimax were purchased from Invitrogen Corporation (Thermo Fisher, Carlsbad, CA). The opti-MEM, M199 culture media, and gentamicin sulfate were purchased from Gibco (Thermo Fisher, Waltham, MA, USA). Collagenase was purchased from Atlanta Biologicals (Flowery Branch, GA, USA). No. 1 glass coverslips, microscope slide, and Chemiluminescent substrate (SuperSignal West Femto) were from Thermo Fisher Scientific (Waltham, MA, USA). Fluoromount-G and clear nail polish were purchased from Electron Microscopy Sciences (H**atfield, PA, USA).** Bovine LH was purchased from Tucker Endocrine Research Institute and forskolin, HDL, and Phorbol-12-Myristate-13-acetate (PMA) were purchased from EMD Millipore (Burlington, MA, USA). Bio-Rad protein assay was purchased from Bio-Rad (Hercules, CA, USA) and the non-fat milk was from local Kroger (Cincinnati, OH, USA). The siRNA, siGLO, and siHSL (ON-TARGETplus Custom siRNA (CTM-385778; HOUSF-000005)) were designed and purchased from Dharmacon (Lafayette, CO, USA). The 8-Br cAMP, Chloroquine diphosphate were purchased from Tocris (Bristol, United Kingdom). TopFluor Cholesterol was purchased from Avanti Polar Lipids (Alabaster, AL, USA). Enzyme-linked immunosorbent assay kit for progesterone was purchased from DRG International, Inc (Springfield, NJ, USA). All antibodies used in the study are found in Table 1.

### Tissue collection, luteal cell preparation, elutriation, and cell culture

Bovine ovaries were collected at a local slaughterhouse from first trimester pregnant cows (fetal crown-rump length < 10 cm). The ovaries were immersed in ice cold 1× PBS and then transported to the laboratory at 4 °C.

Using sterile technique, the corpus luteum was surgically dissected from ovary and cut into sliced using a microtome and surgical scissors. Tissue pieces were dissociated using collagenase (103 U/mL) in basal medium (M199 supplemented with 100 U/ml penicillin G-sodium, 100 µg/ml streptomycin sulfate, and 10 µg/ml gentamicin sulfate) for 45 min in spinning flasks at 37 °C. Following incubation, the supernatant was removed and transferred to a sterile 15 mL culture tube. Cells were then washed 3× with sterile PBS, re-suspended in 10 mL of elutriation media (Calcium-free, DMEM media, 4.0 g/L glucose, 1× Pen Strep, 25 mM HEPES, 0.1 % BSA, and 0.02 mg/mL deoxyribonucleasel; pH 7.2), and placed on ice. Fresh dissociation medium was added to the remaining undigested tissue and incubated with agitation for an additional 45 min. The remaining cells were collected, washed 2× with sterile PBS, and combined with the previous sample. After the final wash, cells were re-suspended in 10 mL of culture medium. Viability of cells was determined using trypan blue and cell concentration was estimated using a hemocytometer prior to cell elutriation.

Freshly dissociated cells were re-suspended in 30 mL DMEM (DMEM media, 4.0 g/L glucose, 1× Penicillin Streptomycin Solution, 25 mM HEPES, 0.1 % BSA, and 0.02 mg/mL deoxyribonucleasel; pH 7.2). For all experiments, dispersed luteal cells were enriched for small luteal cells (SLC) and large luteal cells via centrifugal elutriation as previously described [36]. Cells with a diameter of 15-25 µm were classified as small luteal cells, and cells with diameter > 30 µm were classified as large luteal cells (purity of 50-90% enriched large luteal cells).

### Granulosa cell collection and differentiation

Bovine granulosa cells were isolated from follicles (2-8 mm in diameter) from bovine ovaries collected at local slaughterhouse. Briefly, granulosa cells were collected by scrapping and flushing the inner layers of follicles. Cells were then washed twice in DMEM-F12 and then plated on a 60 mm^2^ culture dish at 5 × 10^6^ cells cells/dish with 10% FBS and maintained at 37 °C in an atmosphere of 95% humidified air and 5% CO_2_, as described above until 80-85% confluency.

Bovine granulosa cells were seeded in 35 mm^2^ culture dishes at 5 × 10^5^ cells cells/dish. Granulosa cells were treated with either 1% FBS or Insulin-Transferrin-Selenium (ITS; 1×) and forskolin (10 µM) and 1% FBS to stimulate the differentiation of GC (dGC) and maintained at 37 °C in an atmosphere of 95% humidified air and 5% CO_2_ for 1 – 7 d. Following incubation, spent media and protein was collected and stored at −20 °C until further analysis.

### Luteal cell preparation and treatments

Enriched small and large luteal cells cultures were plated in 12-well culture dishes at 5 × 10^5^ cells/well and 2 × 10^5^ cells/well, respectively. Cells were cultured in M199 culture media supplemented with 5% FBS, 0.1% BSA and 1× Penicillin Streptomycin Solution, at 37 °C in an atmosphere of 95% humidified air and 5% CO_2_, as described above.

#### Treatment with LH, forskolin, and 8-Br cAMP

Prior to experiment, cells were rinsed with 1× PBS and fresh M199 culture media (0.1% BSA and 1× Penicillin Streptomycin Solution) was placed on cells and equilibrated at 37 °C in atmosphere of 95% air and 5% CO_2_ for 2 h. To determine the effects of LH on activation and stimulation of HSL, cells were treated with media alone, LH (1, 3, 10, 30, or 100 ng/mL), forskolin (10 µM), or 8-Br cAMP (1 mM) for 0, 5, 15, 30, 120 and 240 min, at 37 °C in atmosphere of 95% air and 5% CO_2_. Following incubation cell lysate was collected for western blotting.

#### Treatment with adenoviruses

The adenoviruse expressing β-galactosidase (Ad.βGal) were prepared by Chris Wolford (Ohio State University, Columbus, Ohio) as previously described [37]. The adenoviruses expressing the endogenous green fluorescent protein (Ad.GFP) or the endogenous inhibitor of PKA, Ad.PKI as previously described (19). In brief, enriched small luteal cells were seeded into 12-well culture dishes to determine progesterone concentrations or 6-well dishes for confocal experiments and maintained at 37 °C in an atmosphere of 95% humidified air and 5% CO_2_ for 24 hours prior to adenoviral infection. The Ad.GFP and Ad.PKI or Ad.βGal and Ad.PKI were added to cell culture in serum-free M199 for progesterone and confocal experiments, respectively. After 2 hours, the media was replaced with M199 enriched with 5% FBS and maintained for an additional 24 h at 37 °C in an atmosphere of 95% humidified air and 5% CO_2_. For progesterone analysis, media was changed, and cells were equilibrated for 2 h prior to treatment with control (M199 medium) or LH (100 ng/mL) for 4 h. Following incubation spent media was collected to determine progesterone concentration and cell lysate was collected to determine protein concentration. For confocal experiments media was changed, and cells were equilibrated for 2 h prior to treatment with control (M199 medium), LH (10 ng/mL), 8-Br cAMP (1 mM), for 0 to 6 h. Following incubation cells were prepared for confocal microscopy.

#### Drug inhibition of PKA, HSL, or Lysosome

Cells were pre-treated with H89 (PKA inhibitor; 20, or 50 µM), Pristimerin (0.1, 1, or 10 µM), CAY10499 (HSL inhibitor; 0.1, 1, or 5 µM), or Chloroquine diphosphate (20 mg/mL) for 2 h. Following pre-treatment with appropriate drug inhibitor cells were stimulated with LH (100 ng/mL) for 4 h at 37 °C in atmosphere of 95% air and 5% CO_2_. Spent media and protein were immediately collected and stored at −20 °C until further analysis.

#### siRNA Knockdown of HSL

DNM1L was knocked-down using silencing RNA (siRNA) to determine the effects of DNM1L on progesterone production. In brief, enriched small luteal cell populations were transfected with siGLO (a cy5-labeled non-target siRNA as control) or siHSL (ON-TARGETplus Custom siRNA (CTM-385778; HOUSF-000005)) for 6 h using Lipofectamine RNAimax (Invitrogen Corporation, Carlsbad, CA) in opti-MEM 1 (Gibco; Thermo Fisher, Waltham, MA, USA) culture medium. Following transfection, 5% FBS was added to culture media and maintained for 48-h. Successful knockdown of HSL was also confirmed by western blot for each experiment. Following knockdown of siHSL cells were stimulated with LH (100 ng/mL) for 4 h at 37 °C in atmosphere of 95% air and 5% CO_2_. Spent media and protein were immediately collected and stored at −20 °C until further analysis.

### Isolation of mitochondrial and cytoplasmic fractions

Mitochondria and cytosolic fractions were isolated using a mitochondria isolation kit for cultured cells (Thermo Fisher) per manufactures protocol. Protein in each extracted fraction was determined by the Bio-Rad protein assay (Bio-Rad, Hercules, CA, USA) per manufacturer’s protocol. Samples were resuspended in 6 × Laemmli buffer (60 mM Tris-Cl pH 6.8, 2% SDS, 10% glycerol, 5% β-mercaptoethanol, 0.01% bromophenol blue), placed on a dry heat bath at 100 °C for 6 min, and stored at −20 °C until western blotting analysis.

### Western Blotting Analysis

Following appropriate treatment, cells were immediately placed on ice and rinsed 3 × with 1 mL 1× PBS at 4 °C to remove excess media. Cells were lysed with 50 µL cell lysis buffer (10 mM Tris, 100 mM NaCl, 1 mM EDTA, 1 mM EGTA, 1mM NaF, 20 mM Na4O2O7, 2mM Na3VO4, 1% Triton X-100, 10% Glycerol, 0.1% SDS, and 0.5% Deoxycholate) containing 1× protease and phosphatase inhibitor (Pierce, Thermo Fisher). Cells were removed from culture dish using a cell scraper and the cell lysates were sonicated at 40% power setting (VibraCell, Model CV188) to homogenize cell lysate. Following sonication, cell lysate was centrifuged at 4 °C at 12,000 × *g* for 10 min. Protein in the supernatant was determined by the Bio-Rad protein assay (Bio-Rad, Hercules, CA, USA) per manufacturer’s protocol. Samples were resuspended in 6 × Laemmli buffer and placed on a dry heat bath at 100 °C for 6 min.

Proteins (30 µg/sample) were resolved using 10% SDS-PAGE and then electrophoretically transferred to nitrocellulose membranes. Membranes were blocked with TBS-T (10 mM Tris-HCl pH 7.4, 140 mM NaCl, and 0.1% Tween 20) containing 5% non-fat milk solution at room temperature for 1 h. Membranes were incubated in primary antibody (Table 1) for 24 h at 4 °C for detection of total and phosphorylated proteins. Membranes were rinsed 3 × with 1× TBS-T for 5 min. Membranes were then incubated with appropriate horse radish peroxidase-linked secondary antibody (Table 1) for 1 h at room temperature. Blots were then rinsed 3 × with 1× TBS-T for 5 min each. Chemiluminescent substrate (SuperSignal West Femto; Thermo Fisher Scientific) was applied per manufacturer’s instructions. Blots were visualized using UVP Biospectrum 500 Multi-Spectral imaging system (UVP, Upland, CA, USA) and the percent abundance of immunoreactive protein was determined using densitometry analysis in VisionWorks (UVP).

Total proteins were normalized to ACTB prior to calculation of fold induction. The ratio of phosphorylated HSL to total HSL was determined for each treatment and time point. Fold increases due to treatment (control versus LH) were then calculated.

### Confocal microscopy

Sterile No. 1 glass coverslips (22 × 22 mm) were individually placed in each well of a 6-well culture dish. Enriched small luteal cell cultures were seeded at 5 × 10^5^ cells/well. To determine the effects of LH on colocalization of phospho- and total-HSL with mitochondria and lipid droplets, cells were equilibrated in fresh M199 media enriched with 1% BSA for 2 h prior to treatment with LH (10 ng/mL). Cells were maintained at 37 °C in an atmosphere of 95% humidified air and 5% CO_2_ for 30 min, until termination of experiment.

To determine the effects of LH on colocalization of TopFluor Cholesterol with mitochondrial outer membrane protein, TOM20, cells were pre-treated with 5 µM TopFluor Cholesterol (Avanti Polar Lipids) for 48 h to allow incorporation in lipid droplets. Following incubation, cells were washed 3 × with 1× PBS to remove unincorporated lipid probe from cells. Cells were pre-treated with aminoglutethimide (CYP11A1 inhibitor; 50 µM) for 1 h to prevent cholesterol from exiting the mitochondria as pregnenolone prior to treatment with LH (10 ng/mL) or 8-Br cAMP (1 mM) for 0, 0.5, 1, 2, 4, or 6h. To determine the effects of HSL on LH-induced colocalization of TopFluor Cholesterol with TOM20, cells were pre-treated with aminoglutethimide and CAY10499 (50 µM) for 1 h prior to treatment with LH or 8-Br cAMP. To determine the influence of PKA on LH-induced colocalization of TopFluor Cholesterol with TOM20, cells were transfected with Ad.βGal or Ad.PKI, as described above, prior to treatment with LH or 8-Br cAMP. For all treatment combinations, cells were maintained at 37 °C in an atmosphere of 95% humidified air and 5% CO_2_ until termination of experiment.

Cells were then fixed with 200 µL of 4% paraformaldehyde and incubated at 4 °C for 30 mins. Cells were rinsed 3 × with 1 mL 1× PBS following fixation and then incubated with 200 µL of 0.1% Triton-X in 1× PBS-T (0.1% tween-20) at room temperature for 10 min to permeabilize the membranes. The permeabilized cells were rinsed 3 × with PBS and then blocked in 5% BSA for 24 h at 4 °C. Cells were then rinsed and appropriate antibodies for co-localization (Table 1) were added to each coverslip and incubated at room temperature for 60 min. Following incubation, cells were rinsed 3 × with PBS to remove unbound antibody. Cells were then incubated with appropriate secondary antibodies (Table 1) at room temperature for 60 min. Cells were rinsed 3 × with 1 mL 1× PBS to remove unbound antibody. Following labeling with antibodies, coverslips containing labeled cells were mounted to glass microscope slides using 10 µL Fluoromount-G (Electron Microscopy Sciences). Coverslips were sealed to glass microscope slides using clear nail polish and stored at −22 °C until imaging.

Images were collected using a Zeiss 800 confocal microscope equipped with a 63× oil immersion objective (1.4 N.A) and acquisition image size of 1024 × 1024 pixel (101.31 µm × 101.31 µm). The appropriate filters were used to excite each fluorophore and emission of light was collected between 450 to 1000 nm. Approximately 20 cells were randomly selected from each slide and z-stacked (0.15 µm) images were generated from bottom to top of each cell. The JACoP plug-in was used in Image J software to determine the Manders’ overlap coefficient for each image as previously described [38] and transformed into percent colocalization by multiplying Manders’ overlap coefficient by 100 for all colocalization experiments.

### Progesterone Analysis

Progesterone concentrations from spent media was determined using a commercially available enzyme-linked immunosorbent assay kit (DRG International, Inc, Springfield, NJ, USA) per manufacturer’s protocol.

### Statistical Analysis

Each experiment was performed at least three times each using cell preparations from separate dates of collection and all data are presented as the means ± SEM. The differences in means were analyzed by one-way ANOVA followed by Tukey’s multiple comparison tests to evaluate multiple responses, or by *t*-tests to evaluate paired responses. Two-way ANOVA was used to evaluate repeated measures with Bonferroni posttests to compare means. All statistical analysis was performed using GraphPad Prism software from GraphPad Software, Inc.

## Results

### Expression of components of the steroidogenic machinery progesterone production in the bovine granulosa and luteal cells

Freshly isolated granulosa cells, differentiated granulosa cells, and luteal cells were used to determine the expression of steroidogenic proteins, HSL, NR5A2, STAR, CYP11A1, and 3BHSD (Figure 1A). Steroidogenic proteins, NR5A2, STAR, CYP11A1, and 3BHSD were expressed in differentiated granulosa cells, while HSL was expressed greater in luteal cells (P < 0.05). Furthermore, there was an increase in progesterone production in differentiated granulosa cells on d 2 when compared to undifferentiated cells (P < 0.05). Moreover, this increase in progesterone production in differentiated granulosa cells continued throughout the 7-d time course (P < 0.05; Figure 1B).

**Figure 1.**
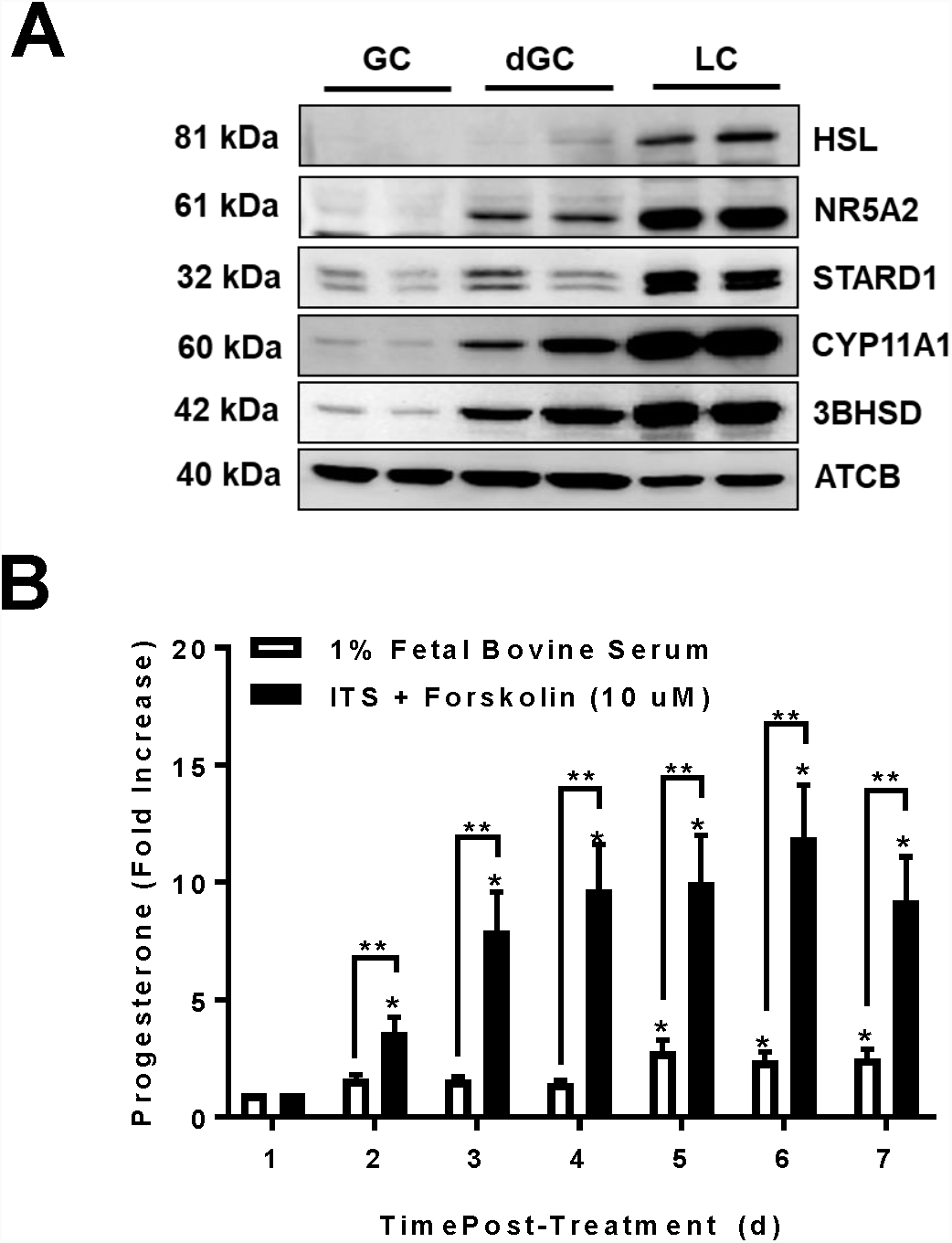
Expression of components of the steroidogenic machinery progesterone production in the bovine granulosa and luteal cells. Freshly isolated bovine granulosa (GC) and luteal cells (LC) were used to determine the expression of steroidogenic proteins. Granulosa cells were treated with Insulin-Transferrin-Selenium (ITS; 1×) and forskolin (10 µM) to stimulate the differentiation of GC (dGC). **Panel A:** Representative western blot analysis of steroidogenic protein expression from bovine granulosa cells (GC), differentiated GC (dGC), and luteal cells (LC). **Panel B:** Progesterone production GC treated with 1% fetal bovine serum (n = 3; open bars) or ITS and forskolin (n = 3; closed bars). Data represented as means ± SEM. *Significant difference within treatment as compared to control at d-1, *P* < 0.05. **Significant difference within day, *P* < 0.05. Hormone sensitive lipase (HSL; 117 kDa); Nuclear Receptor Subfamily 5 Group A Member 2 (NR5A2; 61 kDa); Steroidogenic acute regulatory protein (STARD1; 28 kDa); Cholesterol side-chain cleavage enzyme (CYP11A1; 50 kDa); 3beta-Hydroxysteroid dehydrogenase (3BHSD; 42 kDa); Beta-actin (ACTB; loading control; 45 kDa).

### Effects of luteinizing hormone (LH) on activation and stimulation of hormone sensitive lipase (HSL)

Enriched cultures of small bovine luteal cells were treated for up to 240 min with LH, forskolin, or 8-Br cAMP to determine the influence of LH on stimulation of HSL (Figure 2). Following treatment with LH there was an increase in phospho-HSL at both Ser660 and Ser563 residues. This increase phosphorylation was sustained throughout the experimental period (P < 0.05; Figure 2A). Moreover, treatment with forskolin and 8-Br cAMP stimulated phospho-HSL at both Ser660 and Ser563 residues similar that seen with LH (P < 0.05; Figure 2B and 2C). Additionally, this increase in phospho-HSL at Ser563 was dose-dependent (P < 0.05; Figure 2D).

**Figure 2.**
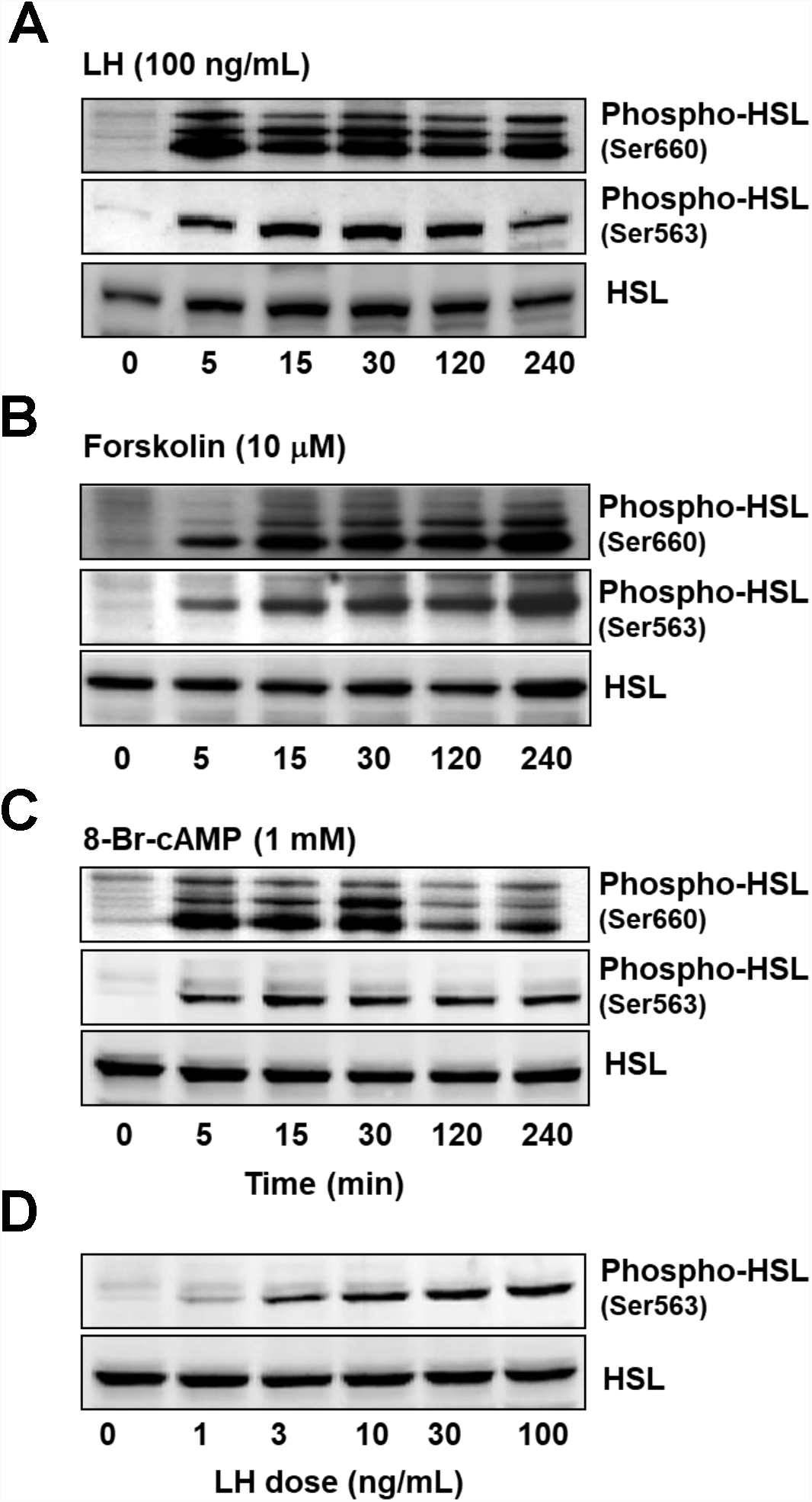
Effects of luteinizing hormone (LH) on activation and stimulation of hormone sensitive lipase (HSL). Enriched cultures of small luteal cells were treated for up to 240 min with luteinizing hormone (LH; 100 ng/mL), forskolin (10 µM), or 8-Br cAMP (1 mM) to determine the influence of LH on stimulation of hormone sensitive lipase (HSL). **Panel A:** Representative western blot analysis for phospho- and total-HSL protein expression in cells treated with LH. **Panel B:** Representative western blot analysis for phospho- and total-HSL protein expression in cells treated with forskolin. **Panel C:** Representative western blot analysis for phospho- and total-HSL protein expression in cells treated with 8-Br cAMP. **Panel D:** Representative western blot analysis for phospho- and total-HSL protein expression in cells treated with increasing concentrations of LH (0-100 ng/mL).

### Effects of Protein Kinase A (PKA) on luteinizing hormone (LH)-induced progesterone production and stimulation of hormone sensitive lipase (HSL)

We utilized a commercially available drug inhibitor for PKA, H89, determine the influence on PKA on activation of HSL and progesterone production in small luteal cells. Pre-treatment with 20 and 50 µM H89 attenuated LH-induced phosphorylation of HSL at Ser563 when compared to control cells (P < 0.05; Figure 3A and 3B). Furthermore, this decrease in phosphor-HSL was accompanied by a decrease in LH-induced progesterone production when compared to control cells (P < 0.05; Figure 3C).

**Figure 3.**
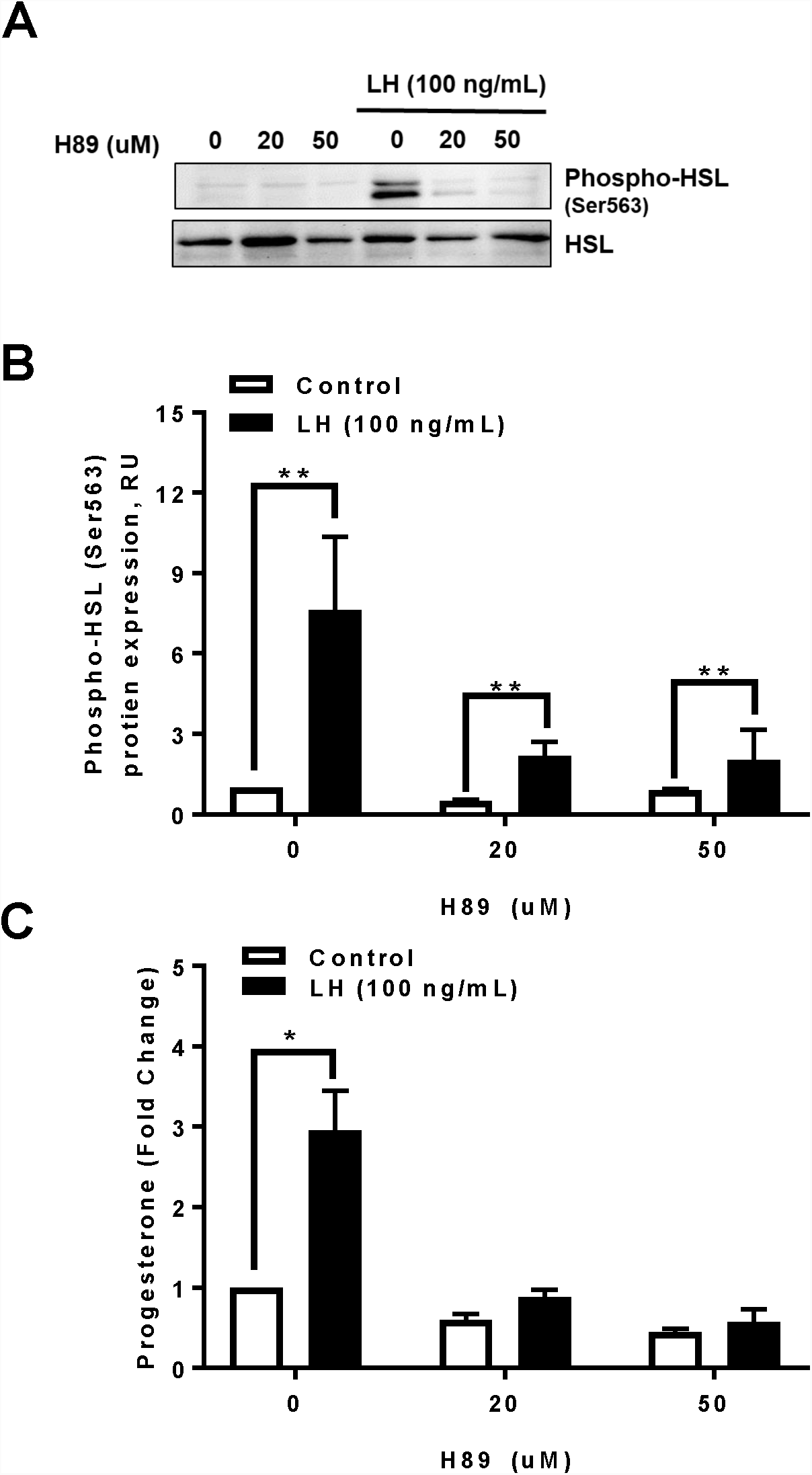
Effects of Protein Kinase A (PKA) on luteinizing hormone (LH)-induced progesterone production and stimulation of hormone sensitive lipase (HSL). Enriched small luteal cells were pre-treated with Protein kinase A (PKA) drug inhibitor H89 and then stimulated with luteinizing hormone (LH; 100 ng/mL) for 4-h. **Panel A:** Representative western blot analysis for phospho- and total-HSL protein expression in cells treated with H89 (Protein Kinase A (PKA) inhibitor; 0,20, or 50 µM) in response stimulation with LH (100 ng/mL). **Panel B:** Densitometric analyses of phospho-HSL (Ser563) protein expression obtained from cells treated with H89 (Protein Kinase A (PKA) inhibitor; 0,20, or 50 µM) in response to 0 ng/mL (n = 3; open bars) or LH (100 ng/mL; n = 3; closed bars). **Panel C:** Progesterone production for enriched small luteal cells pre-treated with increasing concentrations of H89 and stimulated with control (n = 3; open bars) or LH (n = 3; closed bars) for 4 h. Data represented as means ± SEM. **Significant difference within time, *P* < 0.05.

In addition to utilizing H89 to inhibit PKA activity, we used an adenovirus for the endogenous PKA inhibitor, PKI, to determine the effects of PKA on LH-induced progesterone production. In small luteal cells, there a decrease in both basal and LH-induced progesterone production when compared to control Ad.GFP transfected cells (P < 0.05: Figure 4A). Likewise, this attenuation in LH-induced progesterone production was also seen in large luteal cells when compared to control Ad.GFP transfected cells (P <0.05; Figure 4B).

**Figure 4.**
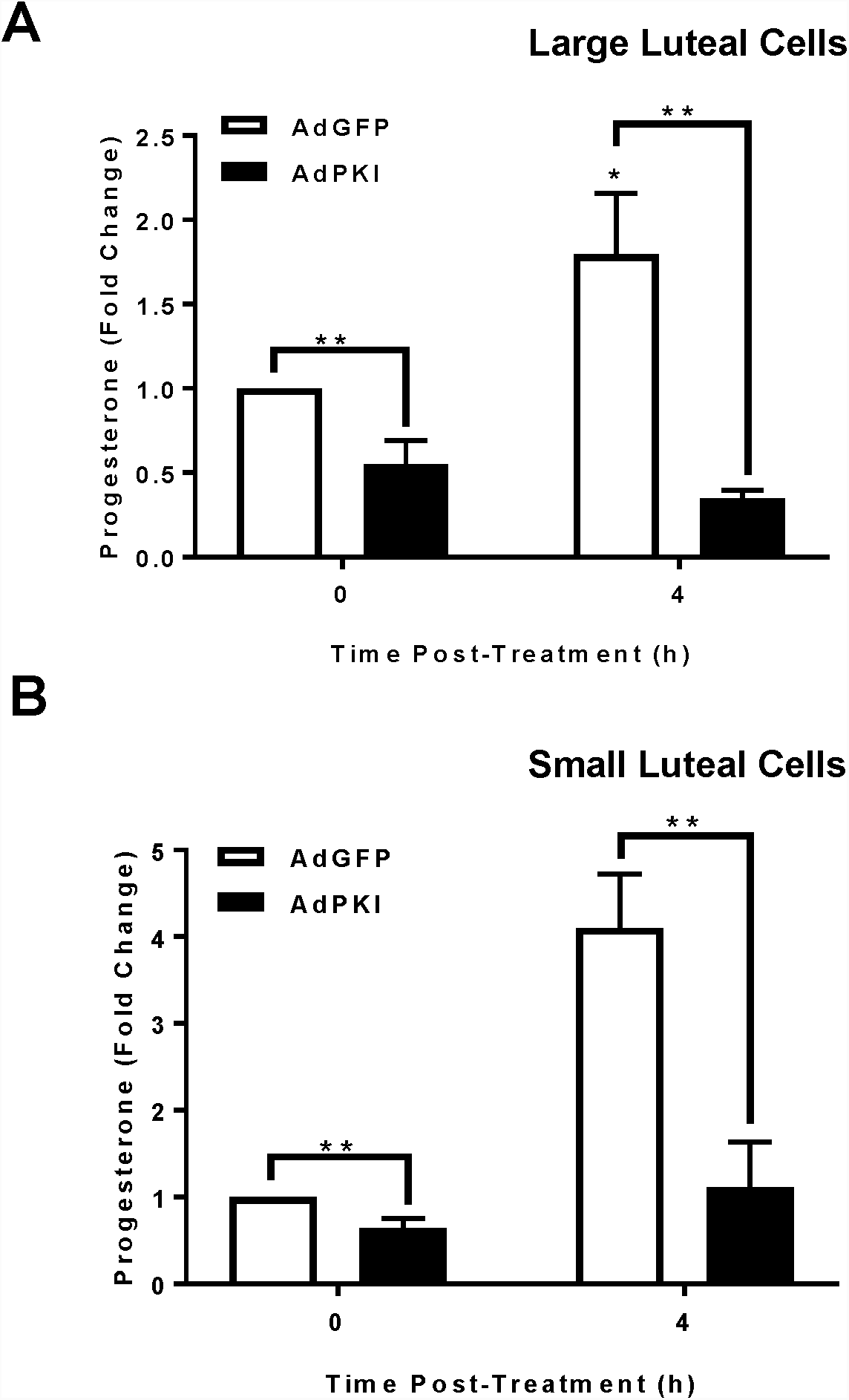
Effects of Protein Kinase A (PKA) on luteinizing hormone (LH)-induced progesterone production in small luteal cells. Replication-deficient adenovirus (Ad) containing the complete sequence of endogenous green fluorescent protein (GFP) cDNA (Ad.GFP) and Protein Kinase Inhibitor (PKI) cDNA (Ad.PKI) was utilized to overexpress GFP or PKI in luteal cells. **Panel A:** Progesterone production for enriched large luteal cells transfected with Ad.GFP (n = 3; open bars) or Ad.PKI (n = 3; closed bars) 4-h post-treatment with luteinizing hormone (LH; 10 ng/mL). **Panel B:** Progesterone production for enriched small luteal cells transfected with Ad.GFP (n = 3; open bars) or Ad.PKI (n = 3; closed bars) 4-h post-treatment with LH. Data represented as means ± SEM. *Significant difference within treatment as compared to control, *P* < 0.05. **Significant difference within time, *P* < 0.05.

### Effects of Hormone sensitive lipase (HSL) on luteinizing hormone (LH)-induced progesterone production

Small luteal cells were treated with Pristimerin or CAY10499 determine the influence of HSL on LH-induced progesterone production. Pre-treatment with 10 µM Pristimerin attenuated LH-induced phosphorylation of HSL at Ser563 when compared to control cells (P < 0.05; Figure 5B). Furthermore, this decrease in phosphor-HSL was accompanied by a decrease in LH-induced progesterone production when compared to control cells (P < 0.05; Figure 5A). Moreover with 5 µM CAY10499 attenuated LH-induced phosphorylation of HSL at Ser563 when compared to control cells (P < 0.05; Figure 5D) and decrease in LH-induced progesterone production when compared to control cells (P < 0.05; Figure 5C).

**Figure 5.**
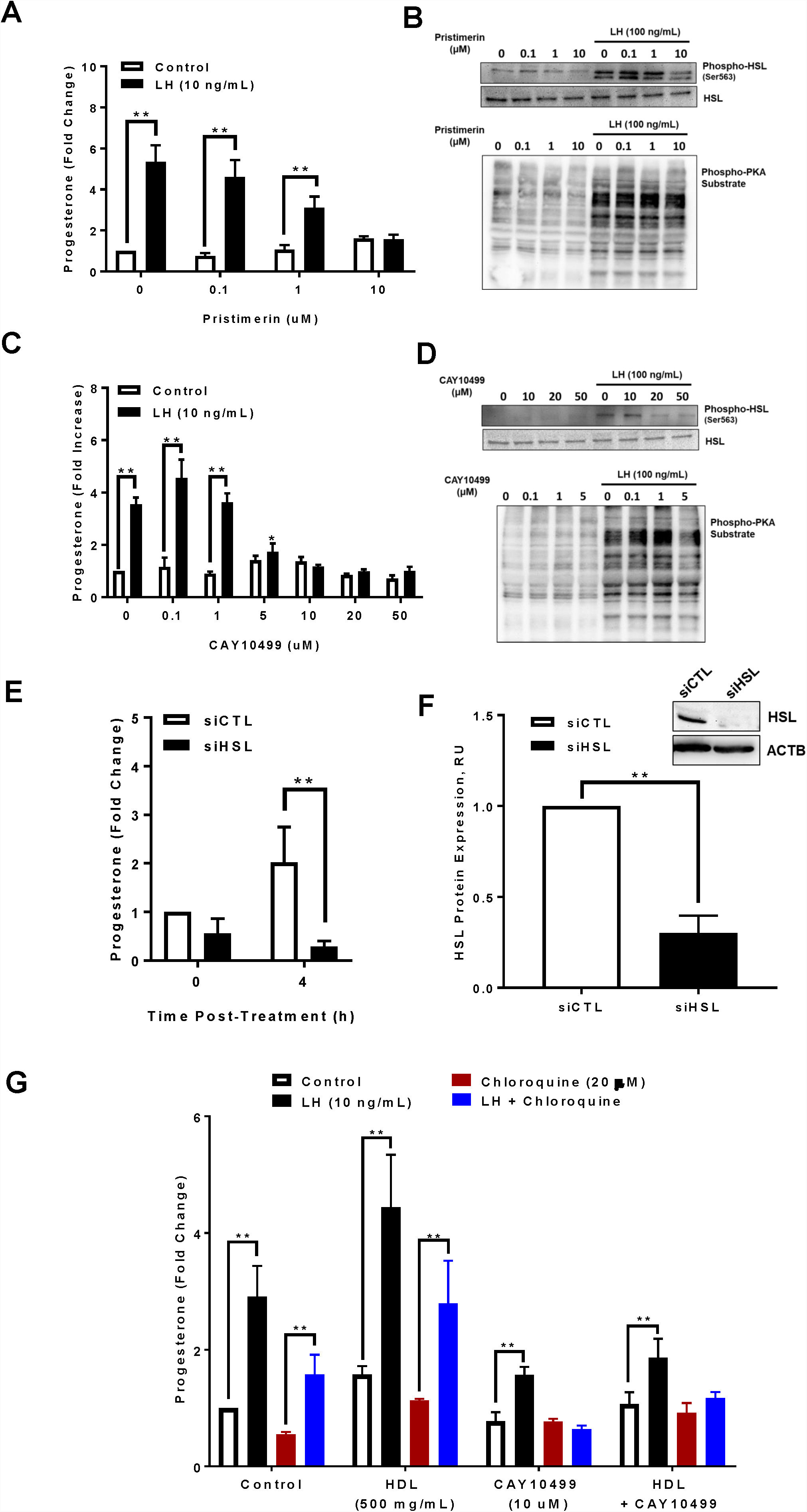
Effects of Hormone sensitive lipase (HSL) on luteinizing hormone (LH)-induced progesterone production. Enriched small bovine luteal cells were pretreated for 1-h with direct and indirect drug inhibitors of hormone sensitive lipase (HSL; Pristimerin, or CAY10499) prior to stimulation with luteinizing hormone (LH; 100 ng/mL). **Panel A:** Progesterone production for enriched small luteal cells treated with increasing concentrations of Pristimerin (0, 0.1, 1, or 10 µM) 4-h post-stimulation with control (n = 3; open bars) or LH (n = 3; closed bars). **Panel B:** Representative western blot analysis for phospho- and total-HSL and phosphor-PKA substrate protein expression in cells treated with increasing concentrations of Pristimerin (0, 0.1, 1, or 10 µM) 4-h post-stimulation with LH. **Panel C:** Progesterone production for enriched small luteal cells treated with increasing concentrations of CAY10499 (0, 0.1, 1, 5, 10, 20, 50 µM) 4-h post-stimulation with control (n = 3; open bars) or LH (n = 3; closed bars). **Panel D:** Representative western blot analysis for phospho- and total-HSL and phosphor-PKA substrate protein expression in cells treated with increasing concentrations CAY10499 (0, 10, 20, 50 µM) 4-h post-stimulation with LH. **Panel E:** HSL mRNA was silenced using siHSL in small luteal cells and stimulated with LH for 4-h. Spent media progesterone obtained from siCTL (n = 3; open bar) and siHSL (n = 3; closed bar) cells 4-h post-treatment with LH. **Panel F:** Representative western blot analysis (insert) showing expression of total-HSL protein expression in siHSL knockdown small luteal cells and densitometric analyses of HSL protein expression obtained from siCTL (n = 3; open bar) and siDNM1L (n = 3; closed bar) validating successful knockdown. **Panel G:** Progesterone production for enriched small luteal cells pre-treated with control, HDL (500 mg/ml), CAY10499 (10 µM), or HDL and CAY10499, and stimulated with control (n = 3; open bars) or LH (n = 3; closed bars), Chloroquine (20 µM; n = 3; red closed bars), or a combination of LH and Chloroquine (n = 3; blue closed bars), for 4-h. Data are represented as means ± standard error. *Significant difference between treatment as compared to control, *P* < 0.05. **Significant difference within treatment, *P* < 0.05.

Another approach to determine the role of HSL employed siRNA targeted for siHSL. Western blotting revealed a 70% decrease in expression of HSL in siHSL-treated luteal cells when compared to control cells (P < 0.05; Figure 5E). Treatment of luteal cells with siHSL resulted in a 44% reduction in basal progesterone secretion. Treatment of luteal cells with siHSL also resulted in a 85% decrease in LH-induced progesterone secretion when compared to control cells, (P < 0.05; Figure 5F).

High density lipoprotein is a source of cholesterol for luteal steroidogenesis. Therefore, to determine if HSL plays a role in HDL processing prior to progesterone biosynthesis, small luteal cells were pre-treated with HDL (500 mg/mL), CAY10499 (10 µM), or a combination of HDL and CAY10499. Cells were then stimulated with LH and/or Chloroquine. Pretreatment with HDL resulted in an increased basal and LH-induced progesterone production when compared to control cells (P < 0.05; Figure 5G). Moreover, in cells pretreated with both HDL and CAY10499, there was a reduction in LH-stimulated progesterone production when compared to control (P < 0.05; Figure 5G) but did not differ from pretreatment of CAY10499 alone (P > 0.05; Figure 5G). Furthermore, pretreating cells with HDL and stimulating with both LH and Chloroquine, lead to an increased in progesterone production when compared to control cells (P < 0.05; Figure 5G) and did not differ from LH stimulation alone (P > 0.05; Figure 5G). Moreover, pretreating cells with HDL and CAY10499 lead a decrease in both LH-induced progesterone production and LH and Chloroquine stimulated progesterone production. Treating small luteal cells with Chloroquine decreased basal progesterone production in control treated cells (P < 0.05; Figure 5G), however in combination with LH, lead to an increased progesterone production when compared to control cells (P < 0.05; Figure 5G) but was decreased when compared to LH alone (P < 0.05; Figure 5G).

### Effects of Luteinizing hormone (LH) on colocalization of phospho- and total-hormone sensitive lipase (HSL) with lipid droplets and mitochondria

Confocal microscopy and western blotting was utilized to determine the effects of LH on localization of phospho- and total-HSL in small luteal cells (Figure 6A and 6B). Here we show an increase in the percent colocalization of both phospho- and total HSL with BODIPY 493/503, following treatment with LH (P < 0.05; Figure 6C). Furthermore, there was an increase in the percent colocalization of both phospho- and total HSL with TOM20, an outer mitochondrial marker, following treatment with LH (P < 0.05; Figure 6D). To validate localization of HSL with lipid droplets and mitochondria, cells were treated with LH, forskolin, and Protein kinase C activator, PMA for 1 h and cytosolic fractions (lipid droplets) and mitochondrial fractions were isolated Here we show localization of both phospho- and total-HSL in both cytosolic and mitochondrial fractions (Figure 6E).

**Figure 6.**
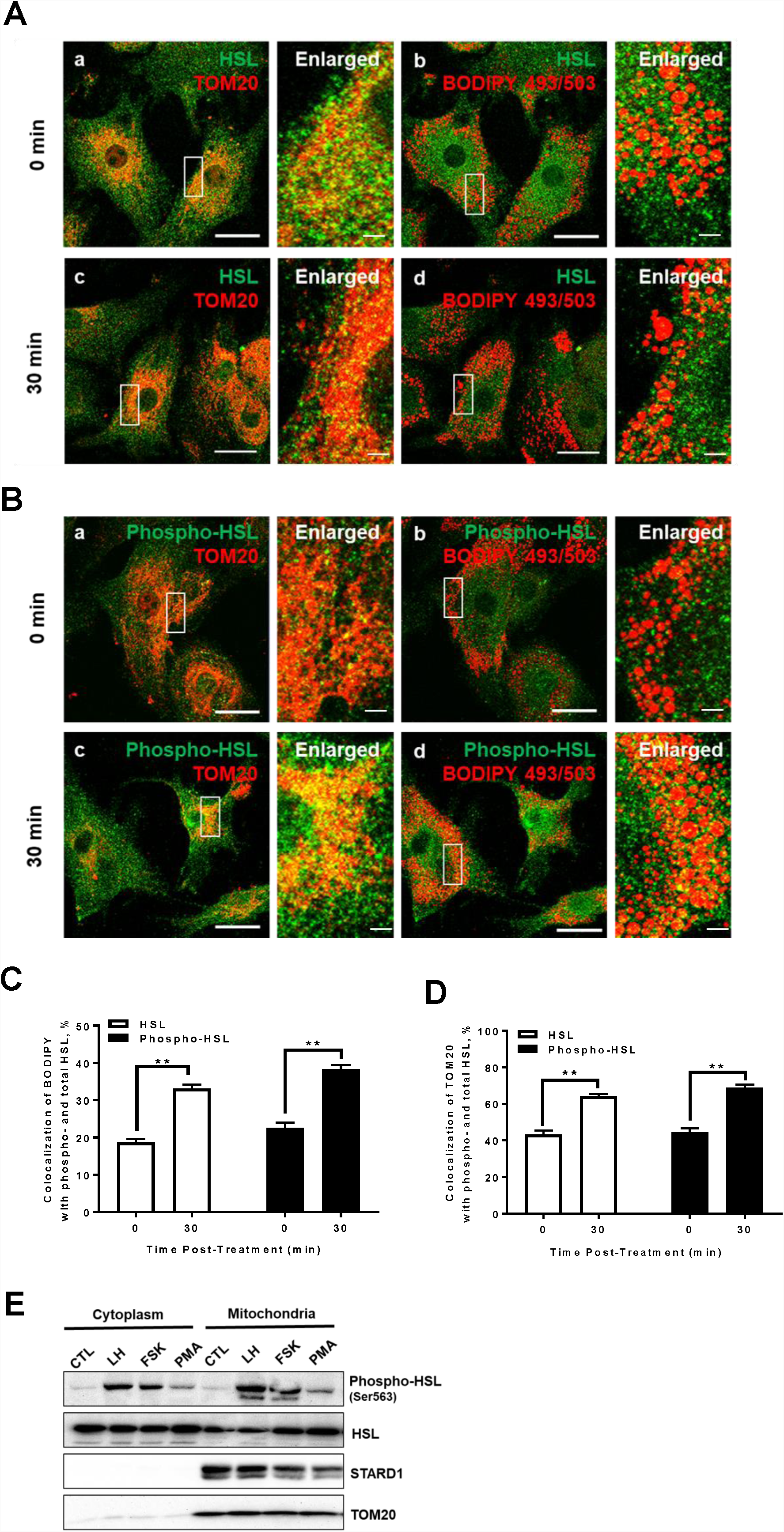
Effects of Luteinizing hormone (LH) on colocalization of phospho- and total-hormone sensitive lipase (HSL) with lipid droplets and mitochondria. Confocal microscopy and western blotting were utilized to determine the effects of LH on localization of phospho- and total-HSL in small luteal cells. **Panel A:** Representative micrographs of total HSL and Tom20 (a and c), total HSL and BODIPY (b and d), and enlarged image (white box corresponding to adjacent image) following stimulation with LH (10 ng/mL). **Panel B:** Representative micrographs of phospho-HSL and Tom20 (a and c), phospho-HSL and BODIPY (b and d) and enlarged image (white box corresponding to adjacent image) following stimulation with LH. **Panel C:** Quantitative analysis of the colocalization of BODIPY with phospho-(n = 3; closed bar) and total-HSL (n = 3; open bar). **Panel D:** Quantitative analysis of the colocalization of TOM20 with phospho-(n = 3; closed bar) and total-HSL (n = 3; open bar). **Panel E:** Representative western blot analysis showing expression of phospho-and total-HSL protein expression in cytoplasmic and mitochondrial cell fractions. Data are represented as means ± standard error. **Significant difference between treatment, *P* < 0.05. Micron bar represents 20 and 2 µm.

### Effects of Luteinizing hormone (LH), Protein kinase A (PKA), and Hormone Sensitive Lipase (HSL) on colocalization of mitochondria with TopFluor Cholesterol in small luteal cells

Confocal microscopy was used to determine whether cholesterol stored in lipid droplets was utilized LH-induced for progesterone biosynthesis. There was an increase in the percent colocalization of TOM20 with TopFluor Cholesterol 4 h following LH stimulation (P < 0.05; Figure 7A, 7C, and Supplementary Figure 2). Moreover, treatment with 8-Br cAMP stimulated an increase in colocalization of TOM20 with TopFluor Cholesterol 4 h following treatment similar that seen with LH (P < 0.05; Figure 7C and Supplementary Figure 3).

**Figure 7:**
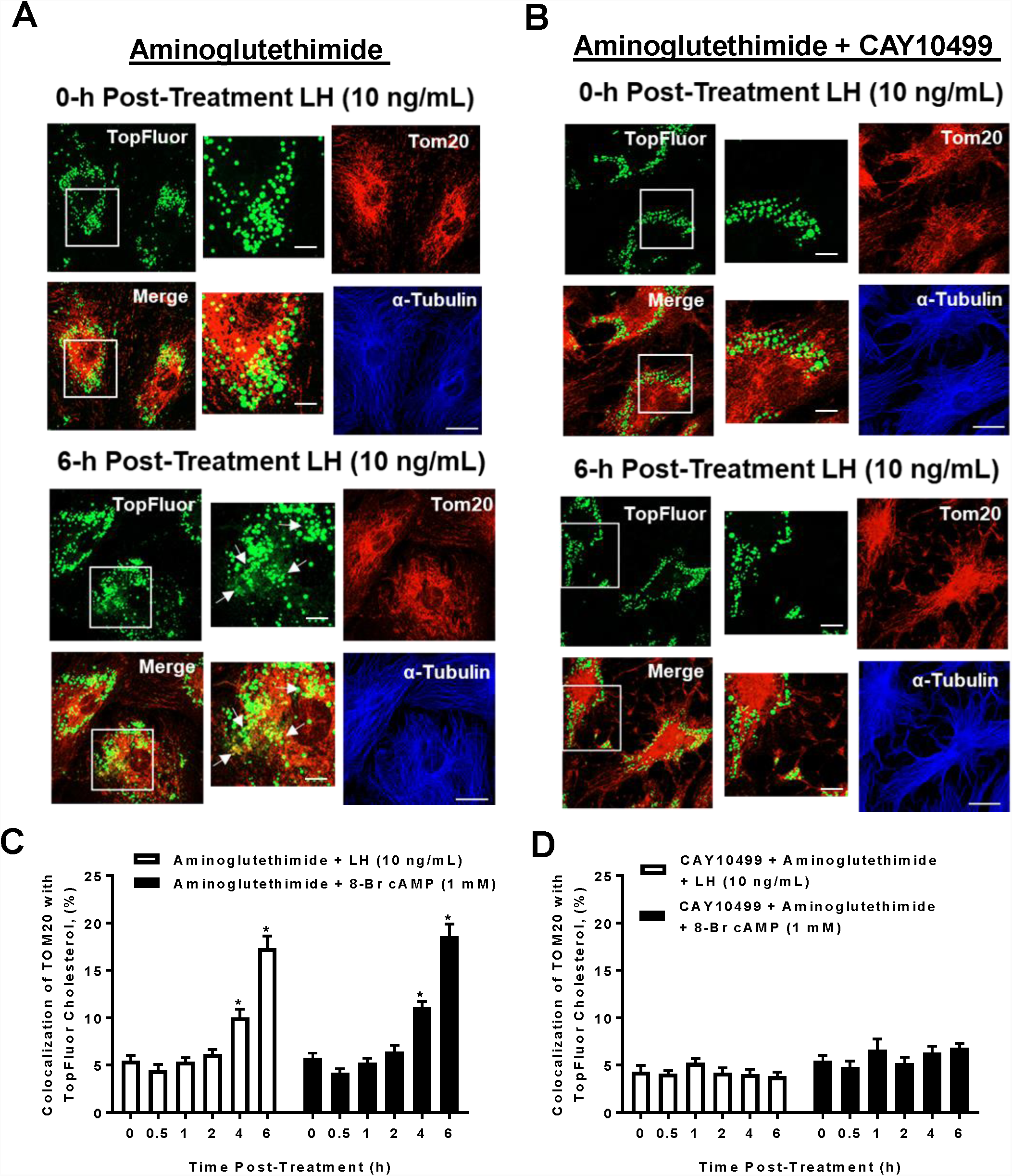
Effects Hormone sensitive lipase on Luteinizing hormone (LH) induced colocalization of TOM20 with TopFluor Cholesterol in small luteal cells. Enriched small luteal cells were preloaded with TopFluor Cholesterol for 48 h. Following cholesterol loading cells were treated with Aminoglutethimide (50 µM) and/or CAY10499 (50 µM) for 1-h prior to stimulation with LH (10 ng/mL) or 8-Br cAMP (1 mM). **Panel A:** Representative micrographs of (left to right) TopFluor Cholesterol, Tom20, merge of TopFluor Cholesterol, and α-Tubulin from cells treated with Aminoglutethimide and stimulated with LH. From top (0-h) to bottom (6-h post-treatment, enlarged image (white box corresponding to adjacent image). **Panel B:** Representative micrographs of (left to right) TopFluor Cholesterol, Tom20, merge of TopFluor Cholesterol, and α-Tubulin from cells treated with Aminoglutethimide, CAY10499 and stimulated with LH. From top (0-h) to bottom (6-h post-treatment, enlarged image (white box corresponding to adjacent image). **Panel C:** Mean colocalization of TOM20 with TopFluor obtained from cells pre-treated with Aminoglutethimide and then stimulated with LH (n = 3 CL; open bar) or 8-Br cAMP (n = 3 CL; solid bar). **Panel D:** Mean colocalization of TOM20 with TopFluor obtained from cells pre-treated with Aminoglutethimide and CAY10499, then stimulated with LH (n = 3 CL; open bar) or 8-Br cAMP (n = 3 CL; solid bar). Data are represented as means ± standard error. *Significant difference between treatment as compared to control, *P* < 0.05. Micron bar represents 20 and 2 µm.

To determine the influence of HSL on LH-induced trafficking of cholesterol from lipid droplets to mitochondria, we inhibited HSL using CAY10499 prior to stimulation with LH. There was no change in the percent colocalization of TOM20 with TopFluor Cholesterol following LH and 8-Br cAMP stimulation for cells treated with CAY10499 (P > 0.05; Figure 7C and Supplementary Figure 4-5).

To determine the influence of PKA on LH-induced trafficking of cholesterol from lipid droplets to mitochondria, we transfected small luteal cells with Ad.βGal (Ad control) or Ad.PKI prior to stimulation with LH. There was an increase in the percent colocalization of TOM20 with TopFluor Cholesterol 4 h following both LH and 8-Br cAMP stimulation in cells transfected with Ad.βGal (P < 0.05; Figure 8A and 8C and Supplementary Figure 6-7). Nonetheless, there was no change in the percent colocalization of TOM20 with TopFluor Cholesterol following LH and 8-Br cAMP stimulation for cells transfected with Ad.PKI when compared to Ad.βGal (P > 0.05; Figure 8C and 8D and Supplementary Figure 8-9).

**Figure 8:**
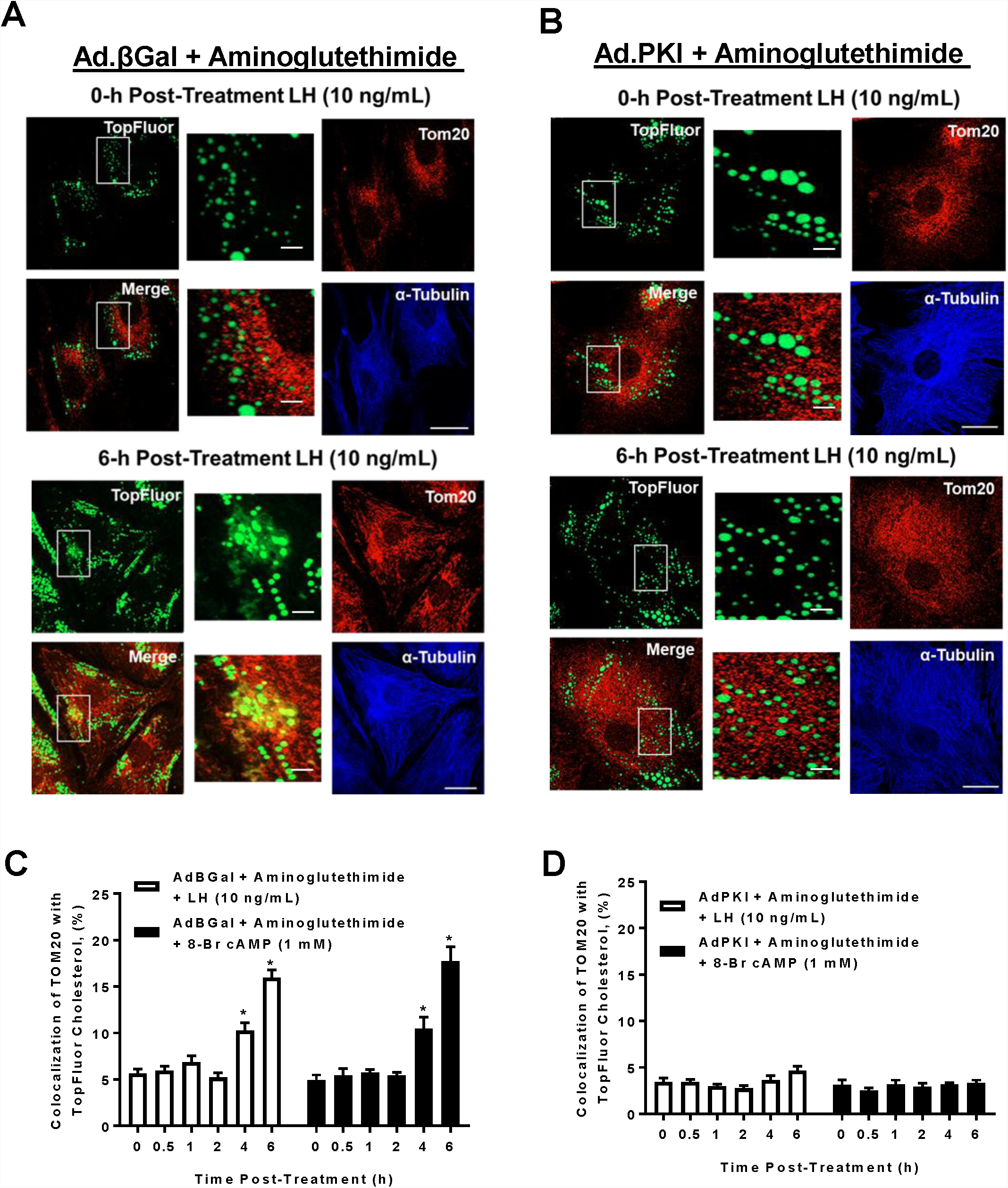
Effects Protein Kinase A (PKA) on Luteinizing hormone (LH) induced colocalization of TOM20 with TopFluor Cholesterol in small luteal cells. Replication-deficient adenovirus (Ad) containing the complete sequence of β-galactosidase (βGal) cDNA (Ad.βGal) and Protein Kinase Inhibitor (PKI) cDNA (Ad.PKI) was utilized to overexpress βGal or PKI in luteal cells. Following transfection, enriched small luteal cells were preloaded with TopFluor Cholesterol for 48 h. Following cholesterol loading cells were treated with Aminoglutethimide (50 µM) for 1-h prior to stimulation with LH (10 ng/mL) or 8-Br cAMP (1 mM). **Panel A:** Representative micrographs of (left to right) TopFluor Cholesterol, Tom20, merge of TopFluor Cholesterol, and α-Tubulin from Ad.βGal transfected cells treated with Aminoglutethimide and stimulated with LH. From top (0-h) to bottom (6-h post-treatment, enlarged image (white box corresponding to adjacent image). **Panel B:** Representative micrographs of (left to right) TopFluor Cholesterol, Tom20, merge of TopFluor Cholesterol, and α-Tubulin from cells treated Ad.PKI transfected cells pre-treated with Aminoglutethimide and then stimulated with LH. From top (0-h) to bottom (6-h post-treatment, enlarged image (white box corresponding to adjacent image). **Panel C:** Mean colocalization of TOM20 with TopFluor obtained from Ad.βGal transfected cells pre-treated with Aminoglutethimide and then stimulated with LH (n = 3 CL; open bar) or 8-Br cAMP (n = 3 CL; solid bar). **Panel D:** Mean colocalization of TOM20 with TopFluor obtained from Ad.PKI transfected cells pre-treated with Aminoglutethimide and then stimulated with LH (n = 3 CL; open bar) or 8-Br cAMP (n = 3 CL; solid bar). Data are represented as means ± standard error. *Significant difference between treatment as compared to control, *P* < 0.05. Micron bar represents 20 and 2 µm.

## Discussion

The present study was undertaken to examine the intracellular signaling events that lead to LH-induced progesterone biosynthesis in luteal cells. Lipid droplets and HSL have been extensively studied in adipose tissue [19, 20, 23, 31], however, the relationship between HSL and lipid droplets is poorly understood in steroidogenic tissues such as the CL. Here we show for the first time HSL acts to drive cAMP/PKA-mediated progesterone biosynthesis in bovine luteal cells following stimulation with LH by initiating hydrolysis of cholesterol esters stored in lipid droplets for acute cholesterol availability.

Lipid droplets have been reported in nearly all cell types, often times as indicators of pathological stress. On the contrary, the presence of lipid droplets in steroidogenic tissues appears to be required for adequate steroid biosynthesis and protection from lipotoxicity. In the ovary, both theca and granulosa cells, contain cytoplasmic lipid droplets. However, in the bovine, luteinization of granulosa cells leads to an accumulation of cytoplasmic lipid droplets as early as day 3 and are presents throughout CL development (D10) [18]. In the rabbit, lipid droplets are increased in luteinized granulosa cells starting at day 1 and are enriched throughout the cytoplasm by day 3 [39]. Similarly, stimulating rhesus macaques with LH or hCG induces lipid droplet-associated PLIN2 mRNA and protein expression in granulosa cells within 12 hours [40]. Here, we show within 2 days of granulosa cells differentiation, there is an increase in progesterone biosynthesis. Moreover, following day 7 of luteinization, there was an increase expression of steroidogenic proteins NR5A2, STAR, CYP11A1, and HSD3B. Additionally, HSL was not detected in differentiated granulosa cells.

It is well-established the luteotropic hormone LH, stimulates adenylyl cyclase leading to increases in intracellular cAMP [7], activation of PKA, and ultimately stimulation of acute steroidogenesis [8, 9]. Yet still, the mechanism by which PKA stimulates optimal steroidogenesis is not fully understood. Here, we show inhibition of PKA leads to both attenuated basal and acute LH-induced progesterone biosynthesis. Moreover, inhibition of PKA impedes the phosphorylation and activation of HSL. In adipocytes, HSL-dependent lipolysis is mediated by PKA-induced phosphorylation, as well as, by the lipid droplet surface protein, PLIN1. Under hormone-activated conditions, active PKA phosphorylates both PLIN1 and HSL. Phosphorylation of HSL leads to the translocation from cytosolic fractions to lipid droplets, specifically interacting with p-PLIN1. In addition, α/β hydrolase domain-containing protein 5 (ABHD5) is released from p-PLIN1 and binds to ATGL to induce lipolysis. In luteal cells, although PLIN3 and PLIN2 are the predominate lipid droplet proteins, a similar cAMP/PKA-dependent mechanism may be utilized to initial lipolysis.

In adipocytes, hormonal stimulation of PKA induced phosphorylation of lipid droplet proteins, perilipin (PLIN) 1 and HSL. Together, these events lead to the mobilization of HSL from cytosolic compartments to the lipid droplets. We show 30 min following stimulation with LH, phospho-HSL is mobilized to both lipid droplets and the outer mitochondria. While localization of active HSL to lipid droplets is in agreement with other studies, it is still unknown if HSL directly interacts with the mitochondria, or if the localization of HSL with outer mitochondria is a result of the intimate relationship between mitochondria and luteal lipid droplets. In steroidogenic cells, STAR is the mitochondrial enzyme responsible for the movement of cholesterol from the outer to the inner mitochondria which is a critical regulatory step in progesterone biosynthesis. In the rat adrenal cortex, active HSL has been shown to directly interact with STAR and facilitates movement of cholesterol from lipid droplets to mitochondria for steroidogenesis [41]. The rational for this interaction is unknown, however, it is thought that STAR may alter the conformations of HSL allowing substrate to access the catalytic site more efficiently. In the current study we show phospho-HSL is mobilized to the mitochondria following stimulation of LH. In the luteal cells, HSL may interact with both the lipid droplets and STAR, promoting optimal steroidogenesis; however, further studies are warranted to better understand the interaction of HSL with the mitochondria.

The large and small luteal steroidogenic cells produce progesterone under both basal and LH-stimulated conditions. However, under hormonal conditions, the acute actions of LH on steroidogenesis require maximum cholesterol availability for optimal progesterone production. Yet, under *in vitro* culture conditions absent of free cholesterol, luteal cells have the capacity to synthesize progesterone. This discrepancy between hormonal and in vitro culture conditions sheds further light on the role of intracellular lipid droplets as significant contributors to luteal steroidogenesis. We have demonstrated both activation of PKA and HSL are required for optimal LH-induced progesterone biosynthesis in small luteal cells. Here, we employed a visual model to further understand the role of lipid droplets in progesterone biosynthesis and functional consequences associated with disruption of PKA/HSL activation. Inhibiting mitochondrial enzyme, CYP11A1, allowed all trafficked labeled cholesterol to remain in the mitochondria for colocalization analysis. We show that following LH stimulation labeled cholesterol initially stored in lipid droplets translocates into the mitochondria for progesterone biosynthesis. Furthermore, inhibition of both PKA or HSL prevents LH-induced colocalization of labeled cholesterol with the mitochondria. These results demonstrate cholesterol esters stored in lipid droplets are utilized for LH-induced progesterone biosynthesis. Likewise, PKA-induced activation of HSL is required for release and trafficking of cholesterol from the lipid droplets to the mitochondria

Cholesterol is the precursor for all steroid hormones. However, unlike the plasma membrane, the mitochondrial membrane is relatively cholesterol poor. Therefore, the demand for a constant supply of cholesterol to the mitochondria is critical for optimal steroid biosynthesis. Lipid droplets serve as neutral lipid reservoirs for steroidogenic tissues, storing cholesterol esters which we demonstrated is utilized for steroid production in luteal cells. Additionally, another source of luteal cholesterol is in the form of HDL which is delivered into the cells through Scavenger receptor class B, type I (SR-BI). To determine if hydrolysis of HDL cholesterol esters employed a HSL-dependent mechanism, small luteal cells were pretreated with HDL and HSL drug inhibitor, CAY10499. Cells pretreated with HDL and CAY10499 had a 58% reduction in LH-induced progesterone when compared to LH-stimulated cells pretreated with HDL alone. Moreover, chloroquine had no influence on progesterone biosynthesis for cells treated with CAY10499, regardless of the presence of HDL or LH, indicating the most likely candidate for the hydrolysis of HDL cholesterol esters in luteal cells is HSL. Taken together, these findings further support a role for a HSL-dependent signaling pathway in response to acute LH stimulation.

Here we have clearly demonstrated that acute LH-induced progesterone biosynthesis requires both cAMP/PKA signaling and activation of HSL in small luteal cells. Moreover, we demonstrate that inhibition of PKA signaling and siRNA knockdown on HSL decreases basal progesterone biosynthesis 35 and 50%, respectively. However, the mechanism of action regulating basal progesterone production in luteinized granulosa cells remains unknown. In the present study, we demonstrate granulosa cells have increased progesterone production 2 days following luteinization; yet, HSL is undetectable. Further studies are warranted to determine the molecular transition occurring in luteinized granulosa cells to luteal cells.

We believe that luteal lipid droplets play a critical role in progesterone biosynthesis by storing cholesteryl esters and interacting with steroidogenic proteins to efficiently produce steroids. While the current study focused on the intracellular signaling events leading to LH-induced progesterone biosynthesis, it is in agreement with other studies demonstrating the role of lipid droplets in steroidogenesis. In a study carried out in mouse model, disruption of lipid droplet structural integrity lead to a 60% reduction in chorionic gonadotropin-induced progesterone production, however, had no influence on the timing of the estrous cycle [42]. This suggest lipid droplets play a key role in acute cholesterol availability. More work is warranted to determine if disruption of luteal droplets will lead to attenuated progesterone biosynthesis. One notable feature observed during luteal development is the formation of lipid droplets, however, the exact mechanism of formation of lipid droplets is still unknown. Studies are warranted to determine the mechanism employing the formation of luteal lipid droplets. In addition, experiments examining the effects LH-stimulation on lipid droplet size, number and composition would greatly benefit the field. Moreover, understanding the impact of diet (over- and undernutrition) on development and maintenance of luteal lipid droplets, lipid composition, and LH-induced steroidogenesis may provide therapeutic insights on infertility.

## ACKNOWLEDGMENTS

Disclosure Statement: The authors have nothing to disclose. The authors thank Janice Taylor and James Talaska at the University of Nebraska Medical Center, Advanced Microscopy Core Facility for their assistance with microscopy. The use of microscope was supported by Center for Cellular Signaling CoBRE-P30GM106397 from the National Health Institutes. This project was supported by Agriculture and Food Research Initiative Competitive Grant no. 2017-67015-26450 from the USDA National Institute of Food and Agriculture; NIH grants R01 HD087402 and R01HD092263, the Department of Veterans Affairs; and The Olson Center for Women’s Health.

## SUPPLEMENTARY FIGURE CAPTIONS

**Figure 1:**
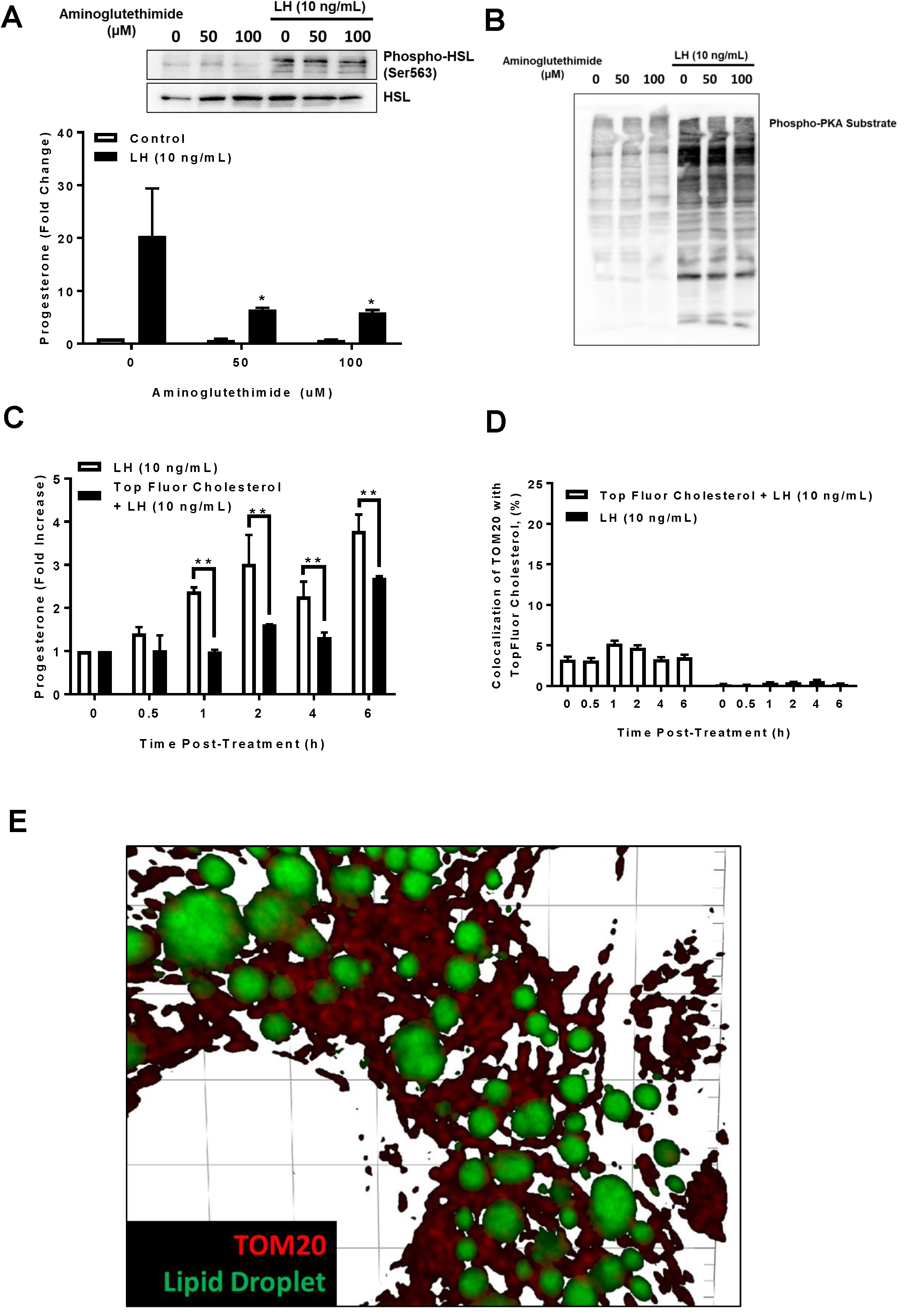
Effects of Aminoglutethimide on Luteinizing Hormone (LH)-induced activation and stimulation of hormone sensitive lipase (HSL) and Protein Kinase A (PKA). Enriched small luteal cells were treated with Aminoglutethimide (0, 50, and 100 µM) and stimulated with LH (10 ng/mL) for 4-h. **Panel A:** Top: Representative western blot of phospho- and total HSL following stimulation with LH (10 ng/mL). Bottom: Progesterone production for enriched small luteal cells pre-treated with Aminoglutethimide (0, 50, and 100 µM) and stimulated with control (n = 3; open bars) or LH (n = 3; closed bars). **Panel B:** Representative western blot of phospho-PKA substrate following stimulation with LH. **Panel C:** Progesterone production for enriched small luteal cells pre-treated with control (n = 3; open bars) or TopFluor Cholesterol (5 µM; n = 3; closed bars) and stimulated LH for 0-6 h. **Panel D:** Mean colocalization of TOM20 with TopFluor Cholesterol (488 nm) obtained from TopFluor negative cells (n = 3 CL; open bar) or TopFluor negative cells (n = 3 CL; solid bar) following stimulation with LH for 0-6 h. **Panel E:** Representative micrograph of TopFluor Cholesterol and TOM20, validating successful pre-loading and gross morphological interactions with mitochondria.

**Figure 2:**
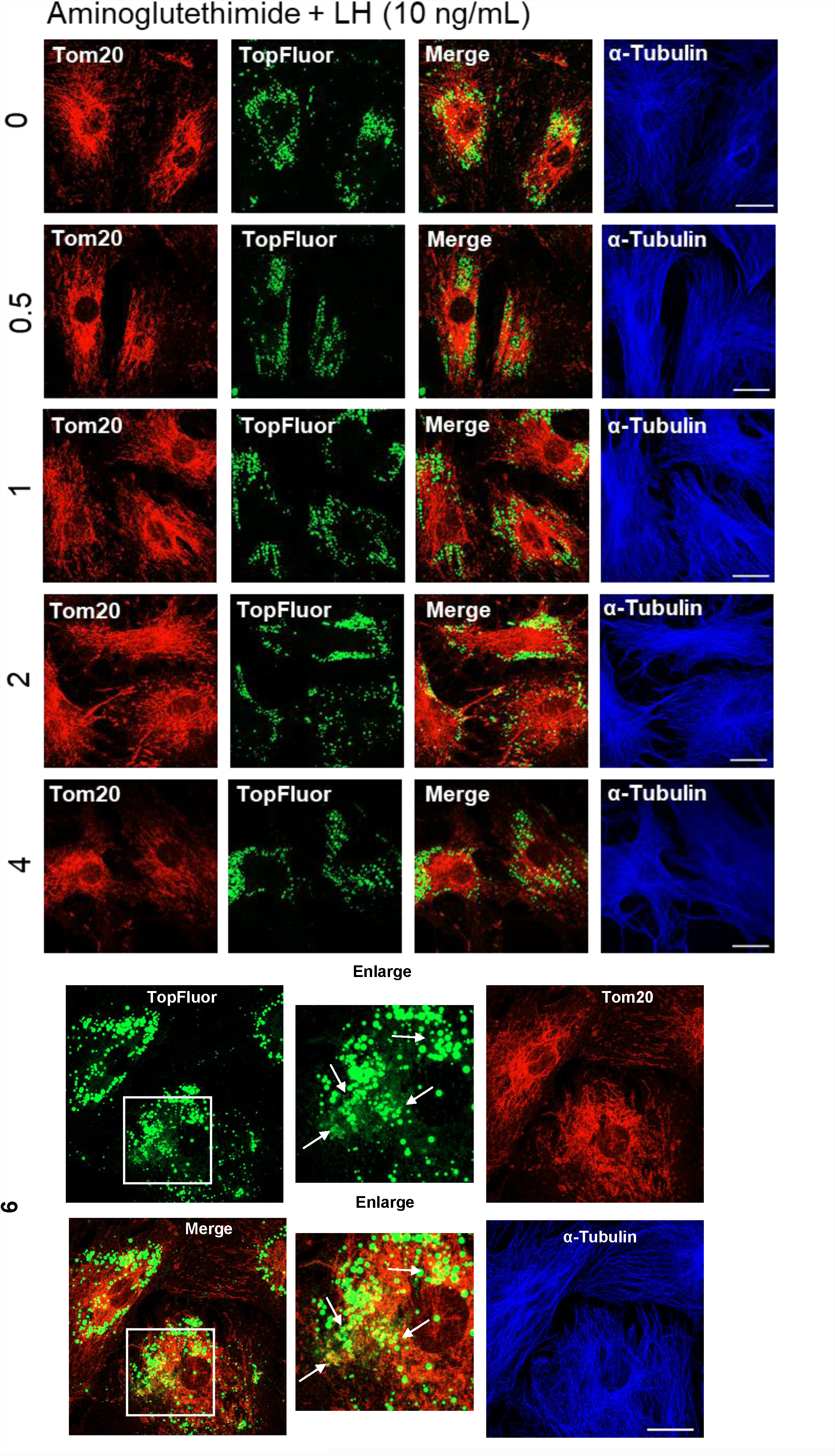
Effects of Luteinizing hormone (LH) on colocalization of TOM20 with TopFluor Cholesterol in small luteal cells. Enriched small luteal cells were preloaded with TopFluor Cholesterol for 48 h. Representative micrographs from cells treated with Aminoglutethimide (50 µM) and LH (10 ng/mL). From top to bottom, time increases from 0 to 6 h. From left to right, Tom 20, TopFluor Cholesterol, merge of Tom20 and TopFluor Cholesterol, and α-Tubulin. Micron bar represents 20 µm.

**Figure 3:**
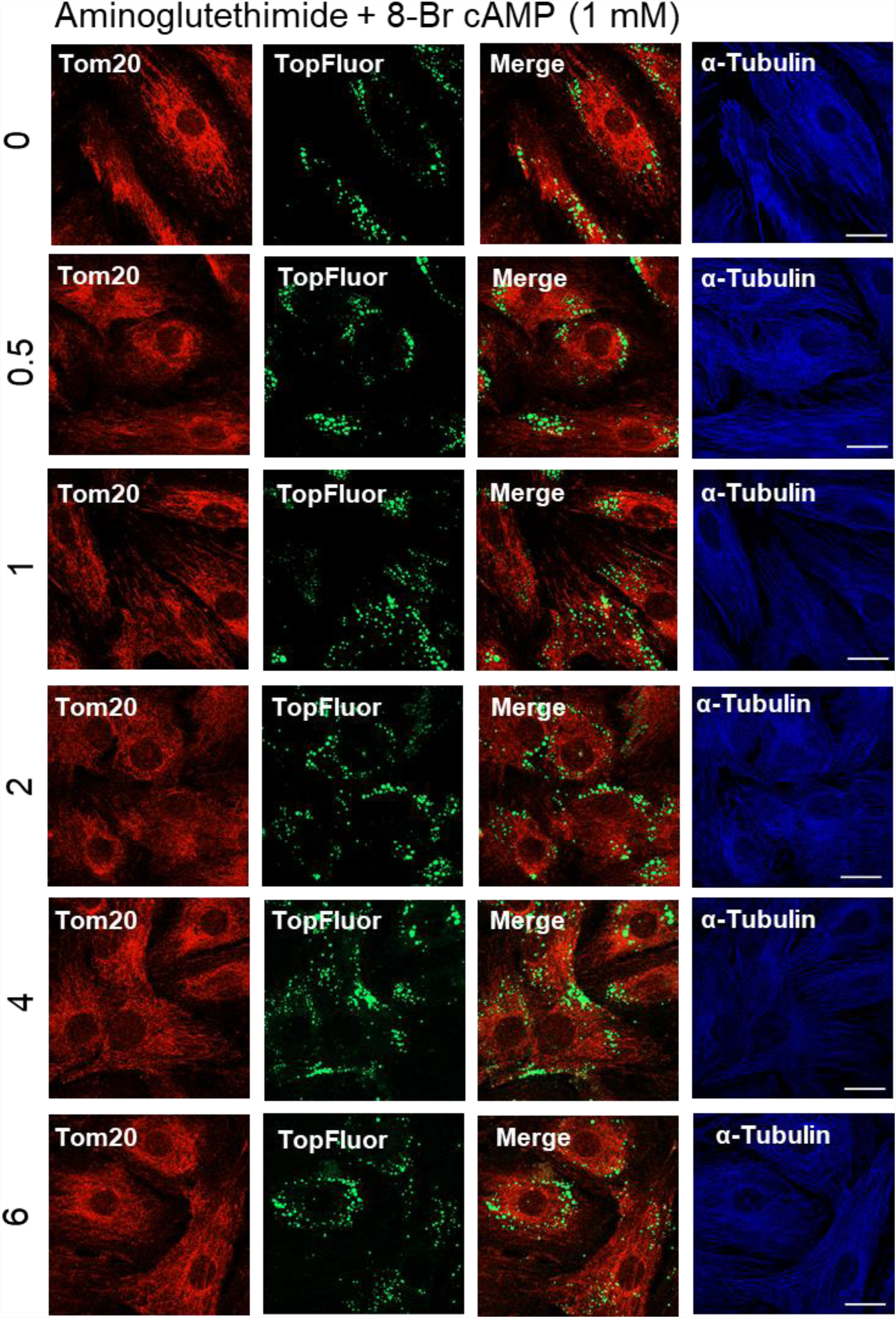
Effects of 8-Br cAMP on colocalization of TOM20 with TopFluor Cholesterol in small luteal cells. Enriched small luteal cells were preloaded with TopFluor Cholesterol for 48 h. Representative micrographs from cells treated with Aminoglutethimide (50 µM) and 8-Br cAMP (1 mM). From top to bottom, time increases from 0 to 6 h. From left to right, Tom 20, TopFluor Cholesterol, merge of Tom20 and TopFluor Cholesterol, and α-Tubulin. Micron bar represents 20 µm.

**Figure 4:**
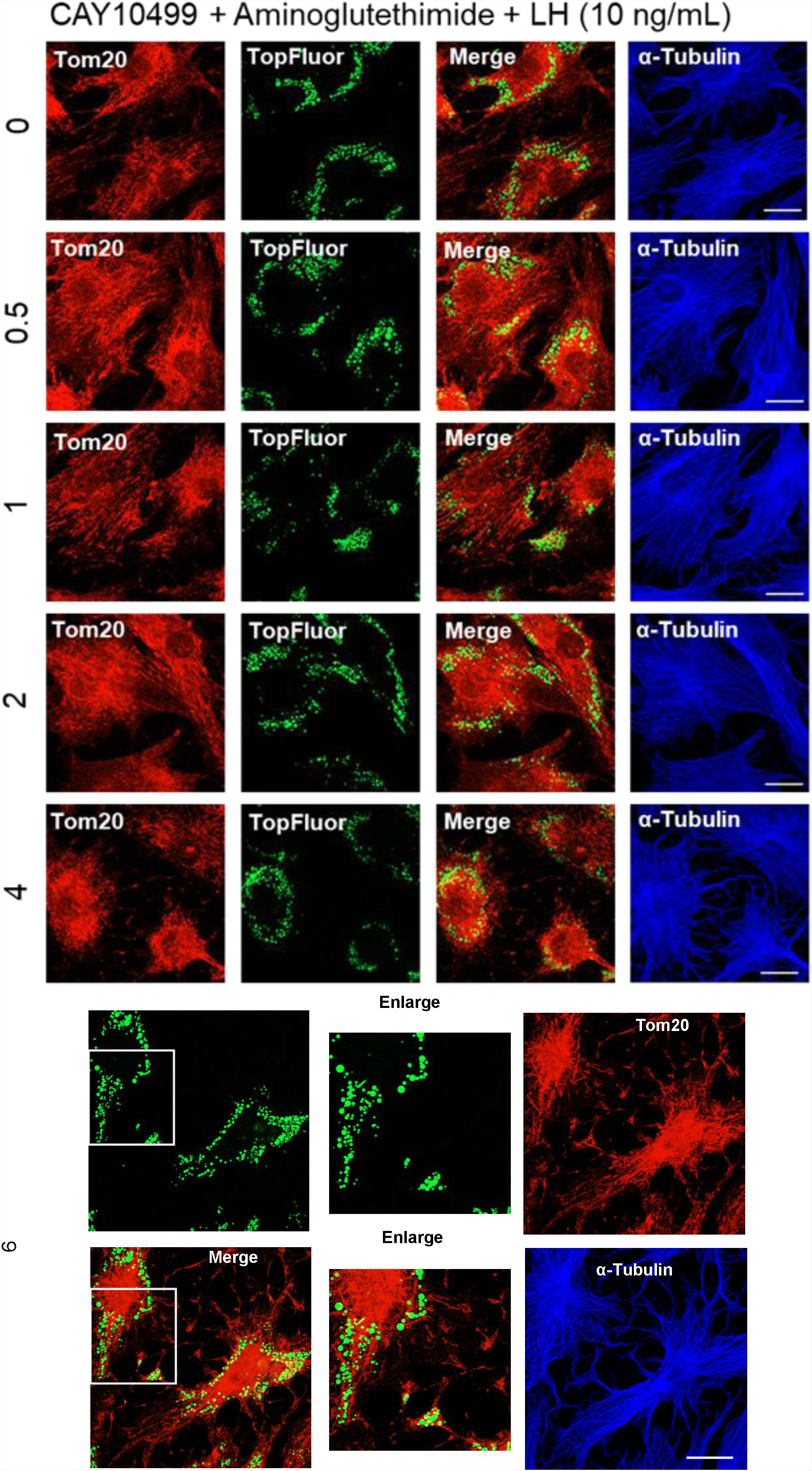
Effects Hormone sensitive lipase on Luteinizing hormone (LH) induced colocalization of TOM20 with TopFluor Cholesterol in small luteal cells. Enriched small cells were preloaded with TopFluor Cholesterol for 48 h. Representative micrographs from cells treated with Aminoglutethimide (50 µM), CAY10499 (50 µm), and LH (10 ng/mL). From top to bottom, time increases from 0 to 6 h. From left to right, Tom 20, TopFluor Cholesterol, merge of Tom20 and TopFluor Cholesterol, and α-Tubulin. Micron bar represents 20 µm.

**Figure 5:**
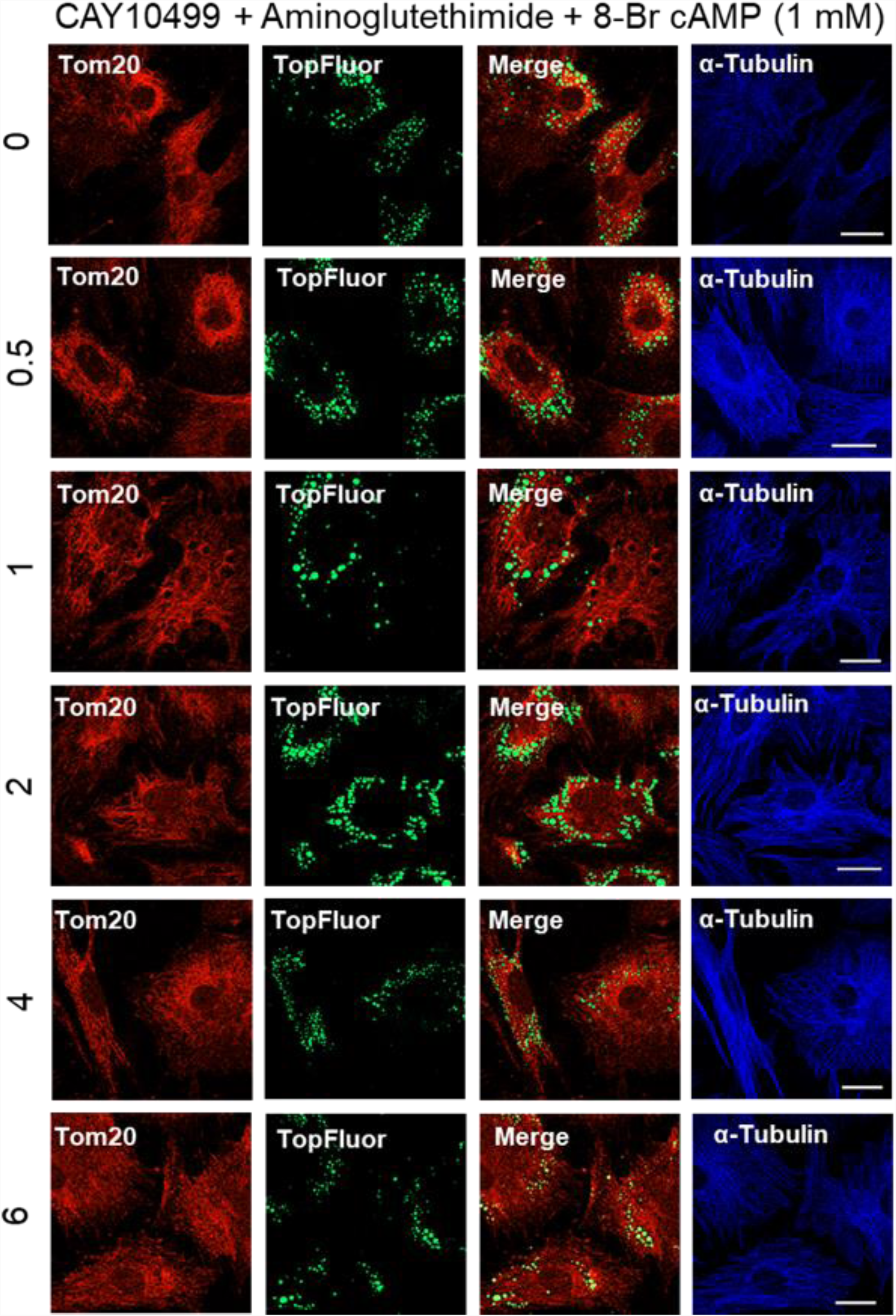
Effects Hormone sensitive lipase on 8-Br cAMP-induced colocalization of TOM20 with TopFluor Cholesterol in small luteal cells. Enriched small cells were preloaded with TopFluor Cholesterol for 48 h. Representative micrographs from cells treated with Aminoglutethimide (50 µM), CAY10499 (50 µm), and 8-Br cAMP (1 mM). From top to bottom, time increases from 0 to 6 h. From left to right, Tom 20, TopFluor Cholesterol, merge of Tom20 and TopFluor Cholesterol, and α-Tubulin. Micron bar represents 20 µm.

**Figure 6:**
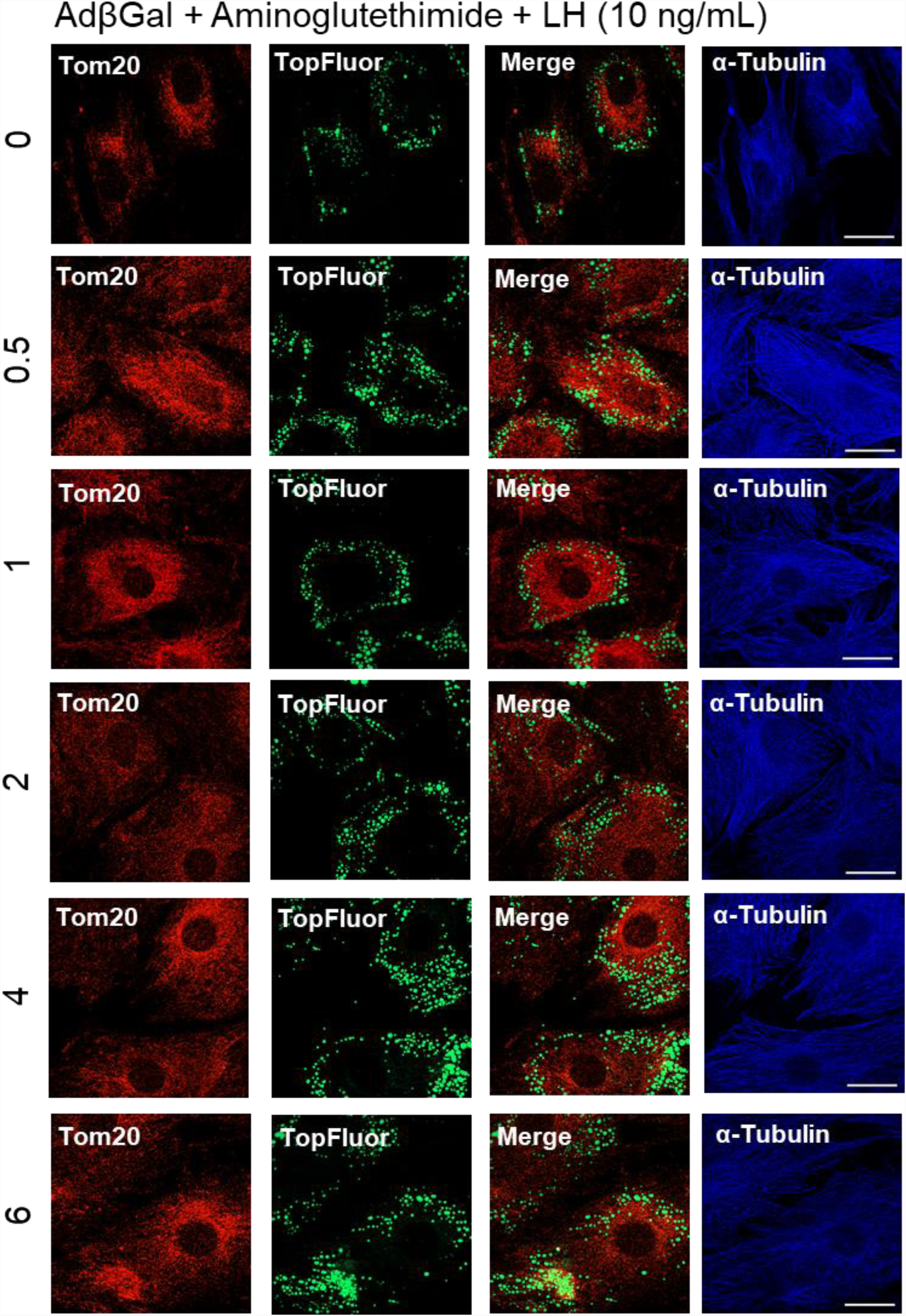
Effects of Luteinizing hormone (LH) on colocalization of TOM20 with TopFluor Cholesterol in small luteal cells transfected with Ad. β**Gal.** Enriched small luteal cells were preloaded with TopFluor Cholesterol for 48 h. Representative micrographs from cells treated with Aminoglutethimide (50 µM) and LH (10 ng/mL). From top to bottom, time increases from 0 to 6 h. From left to right, Tom 20, TopFluor Cholesterol, merge of Tom20 and TopFluor Cholesterol, and α-Tubulin. Micron bar represents 20 µm.

**Figure 7:**
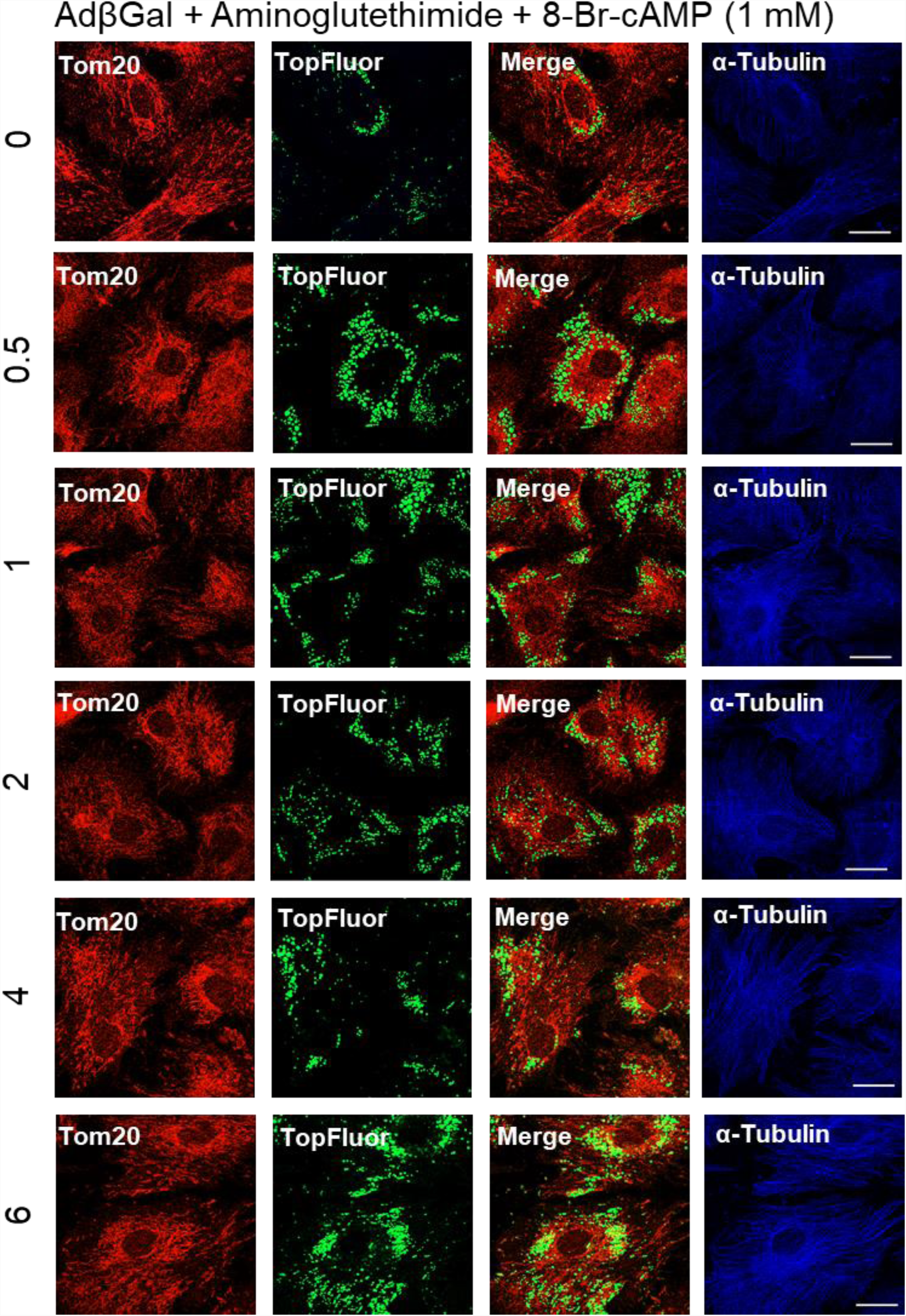
Effects of 8-Br cAMP on colocalization of TOM20 with TopFluor Cholesterol in small luteal cells transfected with Ad. β**Gal.** Enriched small luteal cells were preloaded with TopFluor Cholesterol for 48 h. Representative micrographs from cells treated with Aminoglutethimide (50 µM) and 8-Br cAMP (1 mM). From top to bottom, time increases from 0 to 6 h. From left to right, Tom 20, TopFluor Cholesterol, merge of Tom20 and TopFluor Cholesterol, and α-Tubulin. Micron bar represents 20 µm.

**Figure 8:**
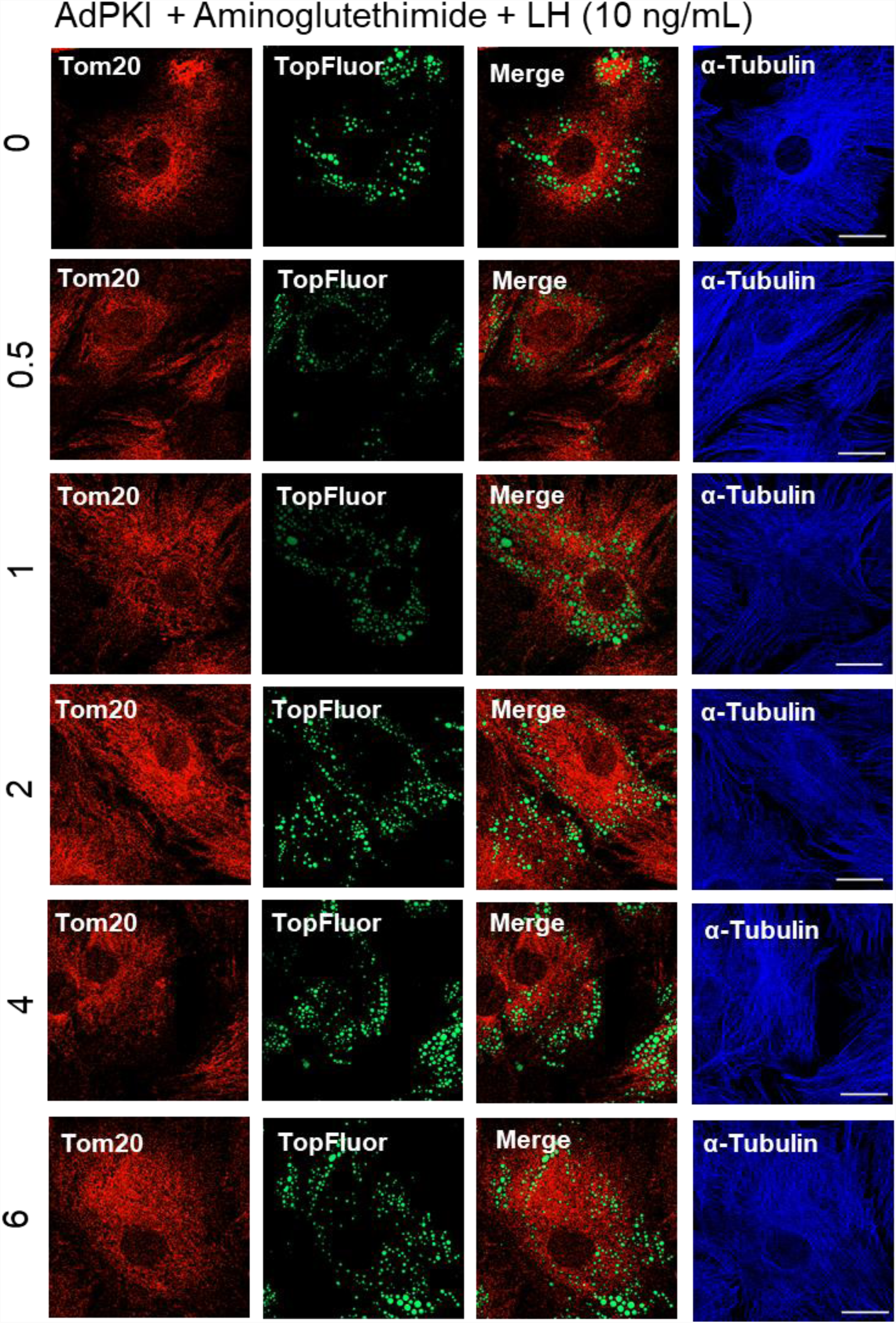
Effects Protein Kinase A (PKA) on Luteinizing hormone (LH)-induced colocalization of TOM20 with TopFluor Cholesterol in small luteal cells. Enriched small luteal cells were preloaded with TopFluor Cholesterol for 48 h. Representative micrographs from cells treated with Aminoglutethimide (50 µM) and LH (10 ng/mL). From top to bottom, time increases from 0 to 6 h. From left to right, Tom 20, TopFluor Cholesterol, merge of Tom20 and TopFluor Cholesterol, and α-Tubulin. Micron bar represents 20 µm.

**Figure 9:**
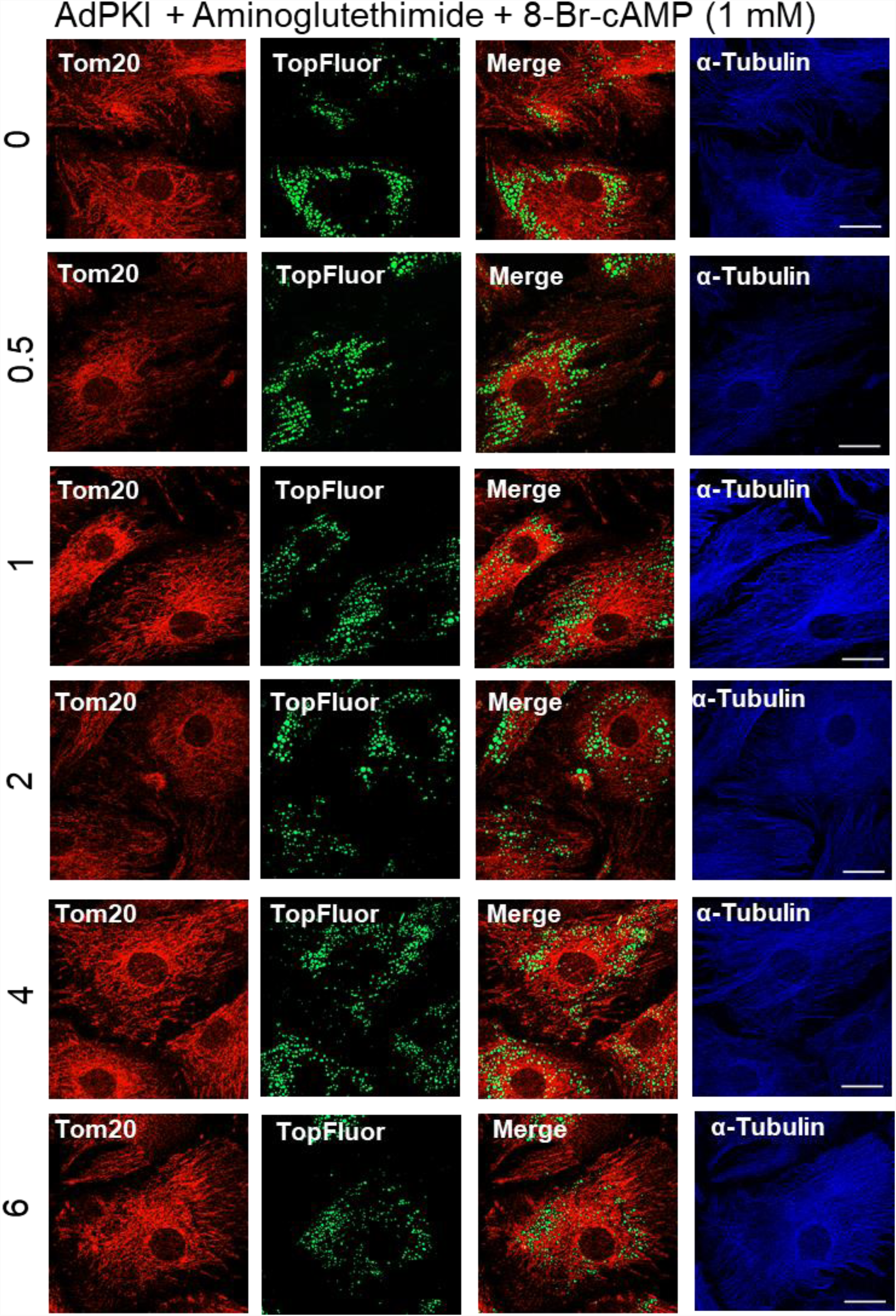
Effects Protein Kinase A (PKA) on 8-Br cAMP-induced colocalization of TOM20 with TopFluor Cholesterol in small luteal cells. Enriched small luteal cells were preloaded with TopFluor Cholesterol for 48 h. Representative micrographs from cells treated with Aminoglutethimide (50 µM) and 8-Br cAMP (1 mM). From top to bottom, time increases from 0 to 6 h. From left to right, Tom 20, TopFluor Cholesterol, merge of Tom20 and TopFluor Cholesterol, and α-Tubulin. Micron bar represents 20 µm.

**Figure 10:**
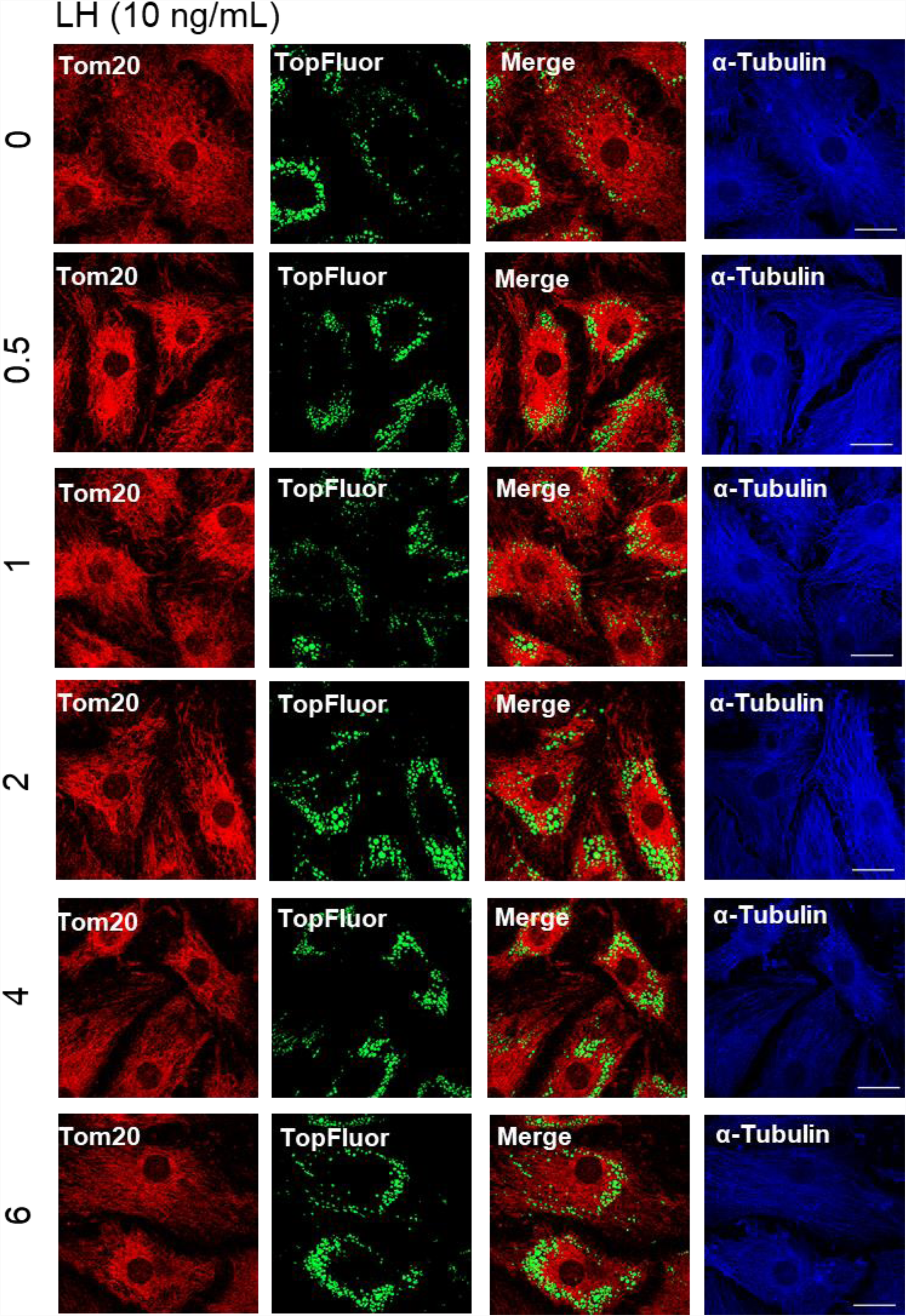
Effects of Luteinizing hormone (LH) on colocalization of TOM20 with TopFluor Cholesterol in small luteal cells. Enriched small luteal cells were preloaded with TopFluor Cholesterol for 48 h. Representative micrographs from cells stimulated with LH (10 ng/mL). From top to bottom, time increases from 0 to 6 h. From left to right, Tom 20, TopFluor Cholesterol, merge of Tom20 and TopFluor Cholesterol, and α-Tubulin. Micron bar represents 20 µm.

**Figure 11:**
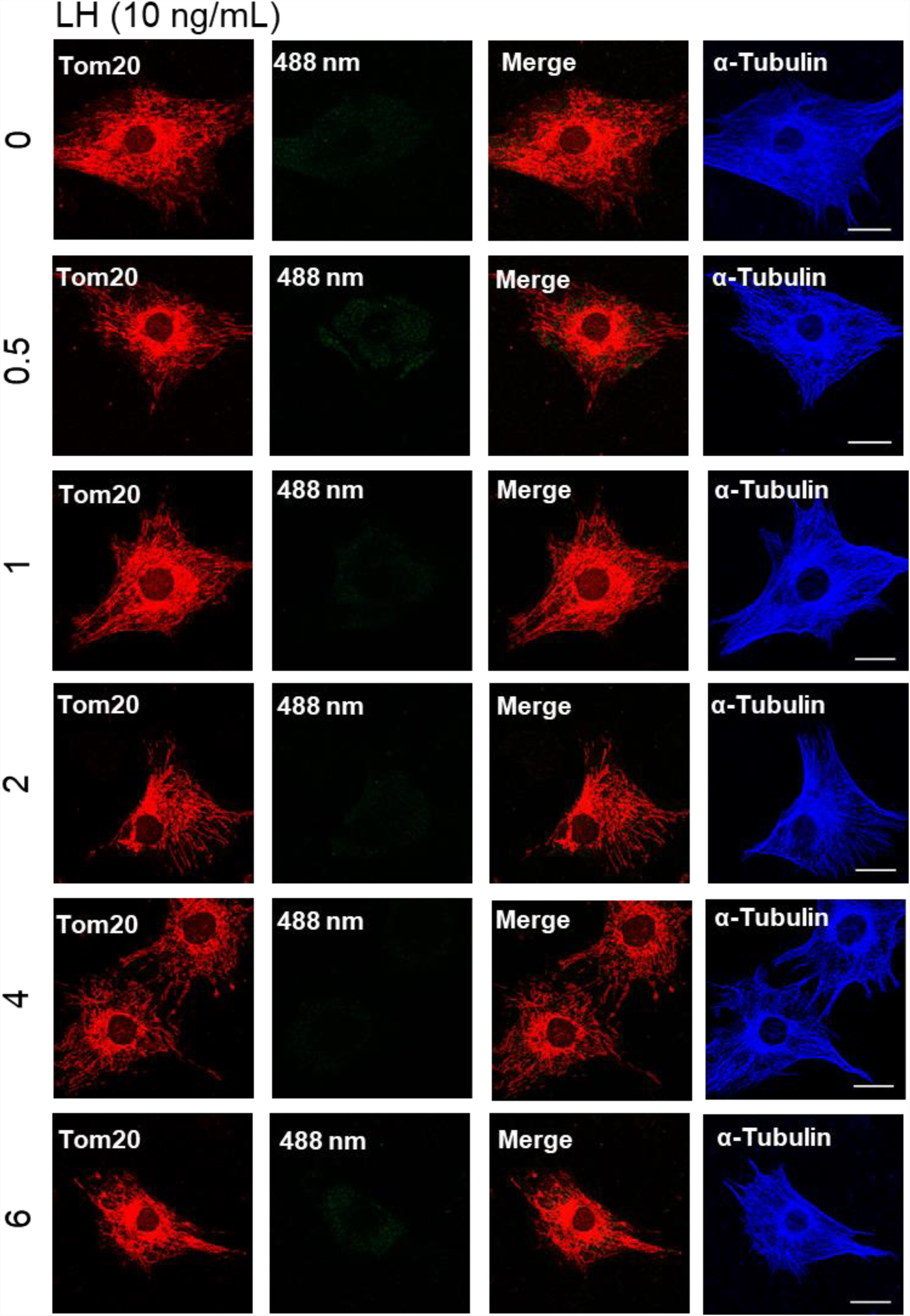
Effects of Luteinizing hormone (LH) on colocalization of TOM20 with TopFluor Cholesterol in small luteal cells. Representative micrographs from cells stimulated with LH (10 ng/mL). From top to bottom, time increases from 0 to 6 h. From left to right, Tom 20, 488 nm channel (neg control), merge of Tom20 and 488 nm channel, and α-Tubulin. Micron bar represents 20 µm.

